# Nucleophagy is promoted by two autophagy receptors and inhibited by chromatin-nuclear envelope tethering in fission yeast

**DOI:** 10.1101/2025.04.09.648038

**Authors:** Zhu-Hui Ma, Zhao-Qian Pan, Zhao-Di Jiang, Guang-Can Shao, Yu Hua, Fang Suo, Chen-Xi Zou, Yi-Feng Jiang, Meng-Qiu Dong, Li-Lin Du

## Abstract

Selective autophagy of the nucleus, known as nucleophagy, targets nuclear components for degradation. The molecular mechanisms underlying nucleophagy remain inadequately understood. In this study, we identify a nucleophagy receptor, Npr1, in the fission yeast *Schizosaccharomyces pombe*. Npr1 is an Atg8-binding multi-transmembrane protein localized to the outer nuclear membrane. It functions redundantly with another autophagy receptor Epr1 to promote nitrogen starvation-induced nucleophagy. In the absence of both Npr1 and Epr1, starved cells exhibit abnormal nuclear morphology and reduced survival. During nucleophagy, the nuclear envelope (NE) forms outward protrusions where Atg8 co-localizes with Npr1 and/or Epr1. These protrusions subsequently detach from the NE, resulting in the formation of autophagosomes that contain nucleophagy cargo. Notably, artificially enhancing chromatin association with the inner nuclear membrane leads to NE protrusions that fail to detach, thereby aborting nucleophagy. Our findings provide mechanistic insights into nucleophagy and suggest that abortive nucleophagy protects chromatin from degradation.

## Introduction

Macroautophagy (hereafter referred to as autophagy) plays a crucial role in maintaining intracellular homeostasis by transporting cytoplasmic components to lysosomes or vacuoles for degradation^1-3^. During autophagy, various cargos, including organelles, are enclosed by an expanding membrane called the phagophore (also known as the isolation membrane), which eventually forms a double-membrane-bound vesicle known as the autophagosome^4,5^. The fusion of the autophagosome with the lysosome or vacuole leads to the degradation of its internal contents^2,6^. Autophagy can be classified into non-selective and selective types^2,3^. Selective autophagy achieves cargo selectivity by employing proteins known as autophagy receptors^7^. Autophagy receptors share a common feature: their ability to interact with the Atg8/LC3 family of proteins through a sequence termed the Atg8-interacting motif (AIM) or the LC3-interacting region (LIR). The core sequence of the AIM/LIR is W/F/YxxL/V/I^8-10^. By simultaneously associating with their cognate cargos and Atg8/LC3 proteins, autophagy receptors establish a physical link between the cargos and the phagophore, thereby promoting selective cargo encapsulation into the autophagosome.

Selective autophagy targets various organelles for degradation, including mitochondria (mitophagy), the endoplasmic reticulum (ER-phagy), and the nucleus (nucleophagy)^11^. The first nucleophagy receptor, Atg39, was discovered in *Saccharomyces cerevisiae* and is found exclusively in budding yeasts^12^. Atg39 is a transmembrane protein localized at the nuclear envelope (NE) and interacts with Atg8 through an AIM in its N-terminal cytosolic tail^12^. In cells undergoing nucleophagy, Atg39 forms bright Atg8-positive puncta in a manner dependent on its Atg8-binding ability^13^. While Atg39 is essential for the selective autophagy of proteins within the outer nuclear membrane (ONM), inner nuclear membrane (INM), nucleolus, and nucleoplasm^12,13^. it is dispensable for the autophagic degradation of nuclear pore components^14-16^. Instead, recent studies have identified the nucleoporin Nup159 as an Atg8-binding protein that promotes the autophagic degradation of nuclear pore components in *S. cerevisiae*^14,15^. The selectivity factors participating in the autophagic degradation of nuclear components in other organisms remain largely undefined.

During nucleophagy in *S. cerevisiae*, histones and DNA are notably absent from nucleophagy-derived autophagosomes^13^. Proteomic analyses also revealed a significant depletion of histones in the contents of autophagic bodies in *S. cerevisiae*^17^. These findings suggest that nucleophagy in *S. cerevisiae* selectively avoids chromatin. However, it remains unclear how this selective exclusion of chromatin is achieved.

In the fission yeast *Schizosaccharomyces pombe*, the soluble autophagy receptor Epr1, which is localized to the ER (including the NE) through its interaction with the integral ER membrane proteins Scs2 and Scs22, mediates ER stress-induced ER-phagy and nucleophagy^18,19^. However, Epr1 is dispensable for nitrogen starvation-induced ER-phagy and nucleophagy^19^. In this study, we identify Npr1, a muti-transmembrane protein localized to the ONM, as a nucleophagy receptor that functions redundantly with Epr1 to facilitate the degradation of nuclear components during nitrogen starvation. Our study also reveals that like the situation in *S. cerevisiae*, the chromatin is excluded from nucleophagy in *S. pombe*. Remarkably, we found that artificially tethering the chromatin to the INM inhibits nucleophagy at the step of NE fission, suggesting a mechanism that prevents chromatin degradation by inhibiting nucleophagy.

## Results

### Nitrogen starvation induces autophagy of specific nuclear components in *S. pombe*

To investigate nucleophagy in *S. pombe*, we examined whether nitrogen starvation triggers the autophagic degradation of specific nuclear components. For this purpose, we analyzed the autophagic processing of five fluorescent protein-tagged markers localized to distinct nuclear subcompartments: Pus1-mECitrine, a nucleoplasmic protein; mECitrine-Bqt4, an inner nuclear membrane protein; mECitrine-Nup82, a nucleoporin; Ker1-mECitrine, a nucleolar protein; and H4-mECitrine, a histone protein associated with chromatin. In wild-type cells, but not in the autophagy-defective *atg5Δ* cells, substantial levels of Pus1-mECitrine **(Fig. 1a)**, mECitrine-Bqt4 **(Fig. 1b)**, mECitrine-Nup82 **(Fig. 1c)**, and Ker1-mECitrine **(Fig. 1d)** were processed into free mECitrine after 24 hours of nitrogen starvation. In contrast, the chromatin marker H4-mECitrine showed no evident processing in wild-type cells **(Fig. 1e)**. These findings indicate that nitrogen starvation in *S. pombe* induces the autophagic degradation of the nucleoplasm, nuclear envelope, nuclear pores, and nucleolus, but not chromatin.

**Fig. 1:**
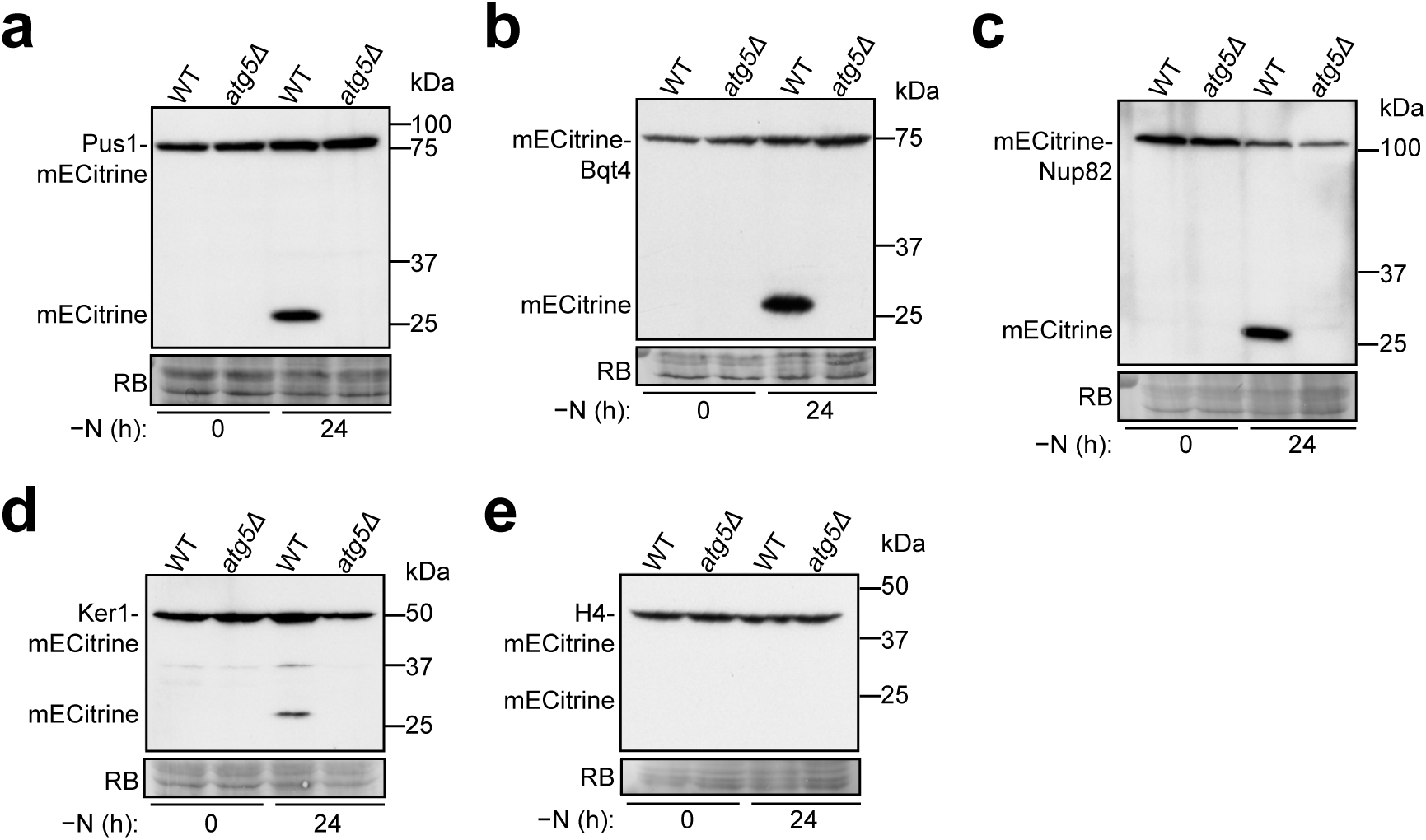
Nitrogen starvation-induced autophagy targets nucleoplasmic, nuclear envelope, nuclear pore, and nucleolar components, but excludes chromatin. Nitrogen starvation-induced autophagic processing of the nucleoplasmic protein Pus1-mECitrine (a), inner nuclear membrane (INM) protein mECitrine-Bqt4 (b), nucleoporin mECitrine-Nup82 (c), nucleolar protein Ker1-mECitrine (d), and histone H4-mECitrine (e) was examined in wild-type and *atg5Δ* cells. Cells expressing mECitrine-tagged proteins from the *P41nmt1* promoter were collected before and after 24 h of nitrogen starvation, and total lysates were analyzed by immunoblotting using an anti-GFP antibody that recognizes mECitrine. Post-immunoblotting staining of the PVDF membrane using Reactive Brown 10 (RB) served as the loading control.

### Identification of Npr1 as a candidate nucleophagy receptor

To identify autophagy receptors involved in nitrogen starvation-induced selective autophagy in *S. pombe*, we employed the TurboID-based proximity labeling technique to search for Atg8-binding proteins^20^. Mass spectrometry analysis of biotin-labeled proteins from nitrogen-starved cells expressing TurboID-tagged Atg8 revealed the enrichment of an uncharacterized protein SPCC70.04c **(Fig. 2a and Supplementary Tabel 1)**, which we named Npr1 (for <u>n</u>ucleo<u>p</u>hagy <u>r</u>eceptor). Co-immunoprecipitation and yeast two-hybrid analyses showed that Npr1 indeed interacts with Atg8 **(Fig. 2b, c)**.

**Fig. 2:**
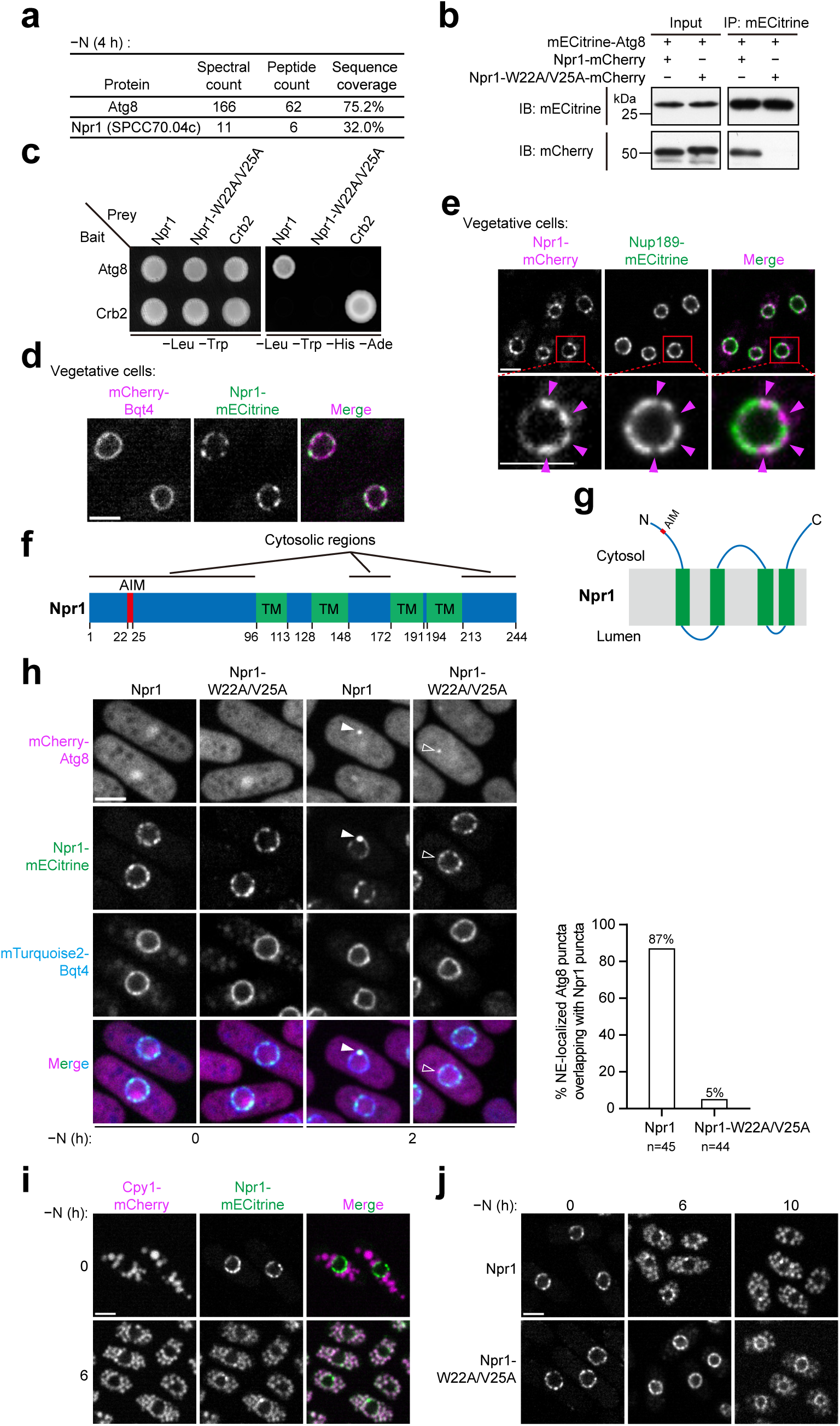
Npr1 is an Atg8-binding nuclear envelope protein that relocalizes to vacuoles during starvation in an Atg8-binding-dependent manner. (a) Proximity labeling identified Npr1 (SPCC70.04c) as an Atg8 interactor. TurboID-tagged Atg8 (expressed under the *P41nmt1* promoter) was used to biotinylate proximal proteins in nitrogen-starved cells. Streptavidin-enriched proteins were analyzed using mass spectrometry. The full mass spectrometry results are presented in Supplementary Tabel 1. (b) Co-immunoprecipitation assays show that Atg8 binds wild-type Npr1 but not the Atg8-interacting motif (AIM)-mutated Npr1 (Npr1-W22A/V25A). (c) Yeast two-hybrid (Y2H) assays confirm that Npr1, but not Npr1-W22A/ V25A, interacts with Atg8. (d) Npr1-mECitrine colocalizes with the NE marker mCherry-Bqt4. Bar, 3 µm. (e) Npr1-mCherry shows a non-uniform NE distribution and tends not to overlap with the nucleopore marker Nup189-mECitrine. A single nucleus is shown in magnified views below, with the positions of Npr1-mCherry signals highlighted by magenta arrowheads. Bar, 3 µm. (f) Membrane topology of Npr1 predicted using CCTOP. Detailed CCTOP output is shown in Supplementary Fig. 2a. Cytosolic regions were validated by split-GFP assays (Supplementary Fig. 2b). TM, transmembrane helices. (g) Schematic of Npr1’s AIM motif and topology (not to scale). (h) AIM-dependent colocalization of Npr1 and Atg8 at the NE during starvation. Left: representative images show that wild-type Npr1, but not AIM-mutated Npr1, forms bright puncta overlapping with NE-localized Atg8 puncta (arrowheads). Bar, 3 µm. Right: quantification of the percentages of NE-localized Atg8 puncta overlapping with Npr1 puncta. (i) Npr1 relocalizes to the vacuole during nitrogen starvation. Cpy1 serves as a vacuole lumen marker. Bar, 3 µm. (j) AIM-mutated Npr1 exhibits a substantial delay in starvation-induced vacuole relocalization compared to wild-type Npr1. Bar, 3 µm.

A proteome-wide study had previously shown that Npr1, when overexpressed, localizes to the nuclear envelope^21^. Consistently, we found that endogenously fluorescent protein-tagged Npr1 exclusively localized to the nuclear envelope in log-phase vegetative cells **(Fig. 2d, e)**. Notably, its distribution on the nuclear envelope was uneven, with a tendency to avoid overlap with nuclear pores **(Fig. 2e)**. The underlying reason for this uneven distribution remains unclear.

Membrane protein prediction analysis indicated that Npr1 is an integral membrane protein containing four transmembrane helices, with both its N- and C-terminal tails facing the cytoplasm or nucleoplasm **(Supplementary Fig. 2a)**^22^. To experimentally assess the membrane topology, we established a split-GFP assay **(Supplementary Fig. 2b)**^23-25^. This assay utilizes two fragments of GFP, GFP_1-10_ and GFP_11_, which can assemble into a fluorescent protein if present in the same cellular compartment. By fusing GFP_1-10_ to proteins of known localization (cytosolic Sum3 or ER luminal Gbs1) and fusing GFP_11_ to either terminus of Npr1, we found that fluorescence was observed only when the Npr1 fusion constructs were co-expressed with the cytosolic reporter, indicating that both termini of Npr1 face the cytosol **(Supplementary Fig. 2b)**. This orientation suggests that Npr1 localizes to the outer nuclear membrane, a notion supported by electron microscopy (EM) analysis using a genetically encoded EM tag, MTn^26^, which showed that MTn-tagged Npr1 was distributed along the outer nuclear membrane **(Supplementary Fig. 2c)**.

To further investigate the interaction between Npr1 and Atg8, we utilized AlphaFold2-Multimer, a tool that effectively predicts binding interfaces between Atg8 and its interacting proteins^27,28^. The predicted structure of the Atg8-Npr1 complex revealed that Npr1 employs a canonical Atg8-interacting motif (AIM), specifically ^22^WIDV^25^, to bind Atg8 **(Supplementary Fig. 2d)**. Mutations of key AIM residues (W22A/V25A) disrupted Npr1’s interaction with Atg8, as demonstrated by both co-immunoprecipitation and yeast two-hybrid analyses **(Fig. 2b, c)**. These findings demonstrate that Npr1 interacts with Atg8 through an AIM located in its N-terminal tail facing the cytosol **(Fig. 2f, g)**.

Under nitrogen starvation conditions, wild-type Npr1, but not its AIM-mutated variant, formed bright puncta that colocalized with Atg8 at the nuclear envelope within 2 hours **(Fig. 2h)**. After 6 hours of starvation, wild-type Npr1 showed pronounced vacuolar re-localization, while the AIM-mutated version exhibited significantly reduced vacuolar targeting **(Fig. 2i, j)**.

Taken together, these results suggest that Npr1 is a candidate nucleophagy receptor. It localizes to the outer nuclear membrane, binds Atg8 via an AIM in its cytosolic N-terminal tail, and undergoes nitrogen starvation-induced vacuolar re-localization in an Atg8-binding-dependent manner.

### Npr1 and Epr1 are redundantly required for starvation-induced nucleophagy

To investigate the role of Npr1 in nucleophagy, we examined the effects of *npr1* deletion on the autophagic processing of various nuclear components during nitrogen starvation. Deletion of *npr1* did not affect the processing of the nucleoplasmic protein Pus1-mECitrine, the inner nuclear membrane protein mECitrine-Bqt4, the nucleolar protein Ker1-mECitrine, or the nucleoporin mECitrine-Nup82 **(Fig. 3a)**. However, overexpression of Npr1 from the *P41nmt1* promoter enhanced the autophagic processing of Pus1-mECitrine, particularly at the early time points of nitrogen starvation, suggesting that Npr1 has a nucleophagy-promoting function **(Supplementary Fig. 3a)**.

**Fig. 3:**
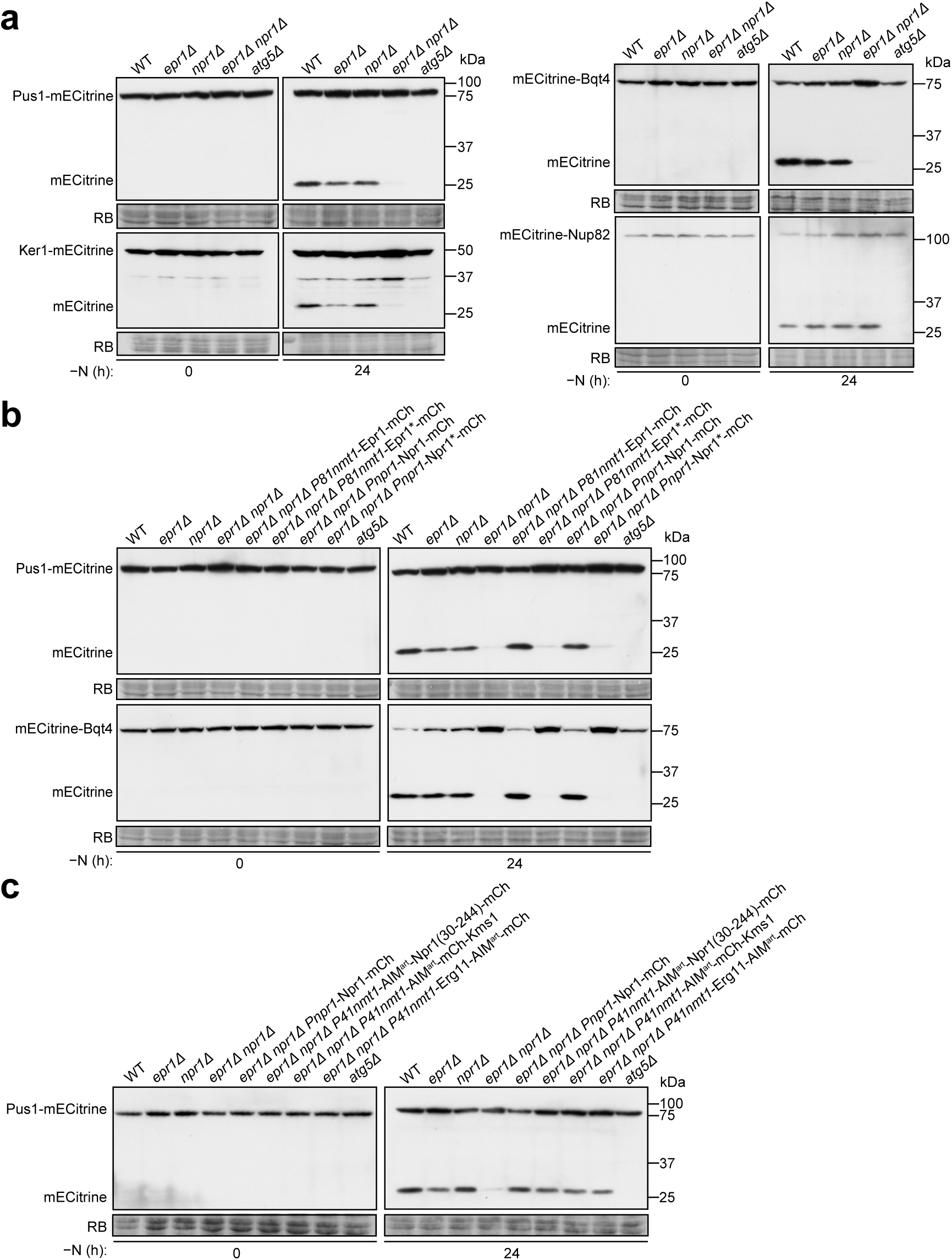
Npr1 and Epr1 are redundantly important for starvation-induced nucleophagy. (a) Nitrogen starvation-induced autophagic processing of the nucleoplasmic protein Pus1-mECitrine, the INM protein mECitrine-Bqt4, and the nucleolar protein Ker1-mECitrine, but not the nucleoporin mECitrine-Nup82, was abolished in *epr1Δ npr1Δ* cells. (b) The nucleophagy function of Npr1 and Epr1 is dependent on their AIMs. Ectopic expression of wild-type Epr1 or Npr1, but not AIM-mutated Epr1 (Epr1*) or Npr1 (Npr1*), in *epr1Δ npr1Δ* cells rescued the nucleophagy defect of *epr1Δ npr1Δ* cells. Epr1*, F352A/V355A; Npr1*, W22A/V25A. (c) Fusing an artificial AIM (AIM^art^), composed of three tandem copies of the EEEWEEL sequence, to either the N-terminally truncated Npr1 lacking its own AIM, or to the cytosol-facing N-terminus of Kms1, or to the cytosol-facing C-terminus of Erg11, rescued the nucleophagy defect of *epr1Δ npr1Δ* cells.

We hypothesized that the absence of a nucleophagy defect in the *npr1Δ* mutant may be due to redundancy with another autophagy receptor. In *S. pombe*, the ER-phagy receptor Epr1 has been shown to be essential for ER-phagy and nucleophagy under ER stress but not during nitrogen starvation^18,19^. Thus, we reasoned that Npr1 and Epr1 may act redundantly under nitrogen starvation conditions. Indeed, deletion of both *npr1* and *epr1*, but not *epr1* alone, nearly completely abolished the nitrogen starvation-induced processing of Pus1-mECitrine, mECitrine-Bqt4, and Ker1-mECitrine **(Fig. 3a)**. Interestingly, the processing of the nucleoporin mECitrine-Nup82 remained unaffected in the *epr1Δ npr1Δ* mutant **(Fig. 3a)**. These results indicate that Npr1 and Epr1 redundantly function as nucleophagy receptors, promoting the autophagic degradation of nucleoplasmic, nuclear envelope, and nucleolar components, but not nuclear pores. This selectivity mirrors the situation in *S. cerevisiae*, where specialized autophagy receptors mediate nucleoporin degradation^14,15^.

Consistent with their redundant roles, re-introducing either Npr1 or Epr1 into the *epr1Δ npr1Δ* mutant fully restored autophagic processing of Pus1-mECitrine and mECitrine-Bqt4 **(Fig. 3b and Supplementary Fig. 3b)**. In contrast, introducing AIM-mutated Npr1 (Npr1-W22A/V25A, abbreviated as Npr1*) or AIM-mutated Epr1 (Epr1-F352A/V355A, abbreviated as Epr1*) failed to rescue the nucleophagy defects **(Fig. 3b and Supplementary Fig. 3b)**, demonstrating that the nucleophagy function of Npr1 and Epr1 depends on their abilities to bind Atg8. Given that Epr1 localizes to both the cortical ER and nuclear envelope^18^, while Npr1 is exclusively localized to the nuclear envelope, we hypothesized that only Epr1 contributes to cortical ER-phagy. Consistent with this idea, nitrogen starvation-induced autophagic processing of the cortical ER membrane protein Rtn1-mECitrine was moderately reduced in the *epr1Δ* mutant but was unaffected in the *npr1Δ* mutant **(Supplementary Fig. 3c, d)**. Furthermore, the additional deletion of *npr1* did not exacerbate the mild phenotype observed in the *epr1Δ* mutant **(Supplementary Fig. 3c, d)**. These results suggest that other, yet unidentified, ER-phagy receptors may act redundantly with Epr1 during nitrogen starvation-induced cortical ER-phagy.

We previously showed that an artificial AIM (AIM^art^) fused to the ER membrane protein Erg11 could functionally substitute for Epr1 in ER stress-induced ER-phagy^18^. To determine whether the nucleophagy functions of Npr1 and Epr1 could similarly be replaced, we employed the same AIM^art^, composed of three tandem copies of the EEEWEEL sequence^29,30^. Substituting the N-terminal 29 amino acids of Npr1 (which contains its AIM) with AIM^art^ generated a chimeric protein, AIM^art^-Npr1(30-244), which successfully rescued the nucleophagy defect of the *epr1Δ npr1Δ* mutant **(Fig. 3c)**, indicating that AIM^art^ can functionally substitute for the AIM of Npr1.

We next fused AIM^art^ to various integral membrane proteins present on the NE. Only the ONM protein Kms1 and the ER membrane protein Erg11 with cytosol-facing AIM^art^ fusions rescued the nucleophagy defect of *epr1Δ npr1Δ* **(Fig. 3c and Supplementary Fig. 3e)**. In contrast, fusion of AIM^art^ to the lumen-facing tails of these proteins, or to the nucleoplasm-facing N-terminus of the INM protein Man1 or the cytosol-facing C-terminus of the cortical ER membrane protein Rtn1, failed to rescue the defect. These findings indicate that the nucleophagy roles of Npr1 and Epr1 can be substituted by a membrane protein localized at the ONM, provided it possesses an AIM that faces the cytosol.

Taken together, these results demonstrate that Npr1 and Epr1 redundantly promote nitrogen starvation-induced nucleophagy by establishing a physical link between the ONM and Atg8. This link is mediated by their AIMs, which are essential for their function.

### Spatiotemporal analysis of Atg8 and nucleophagy receptors at the NE

To further investigate the mechanisms of nucleophagy, we used live-cell imaging to study the genetic interdependency, co-localization, and temporal dynamics of Atg8, Npr1, and Epr1. First, we observed that Atg8 puncta still formed at the NE in *epr1Δ npr1Δ* cells, albeit at slightly lower levels than in wild-type cells **(Supplementary Fig. 4a, b)**. Conversely, NE-localized Npr1 and Epr1 puncta were completely absent in *atg8Δ* cells, indicating that their formation strictly relies on Atg8 **(Supplementary Fig. 4c, d)**.

Next, we analyzed the co-localization patterns of Atg8, Npr1, and Epr1 at the NE using time-lapse imaging of wild-type cells **(Fig. 4)**. We classified NE-localized Atg8 puncta into four types based on their overlap with Npr1 and Epr1 for at least one time point: Type I (36%), which overlapped with both; Type II (55%), with only Npr1; Type III (8%), with only Epr1; and Type IV (1%), with neither **(Fig. 4a, e and Supplementary Fig. 4e, f)**. Thus, in wild-type cells, nearly all NE-localized Atg8 puncta overlapped with receptor puncta, suggesting that they represent nucleophagy events. The fact that over half of the NE-localized Atg8 puncta overlapped with only one receptor indicates that Npr1 and Epr1 can function independently, consistent with genetic data showing that either receptor is sufficient for nucleophagy.

**Fig. 4:**
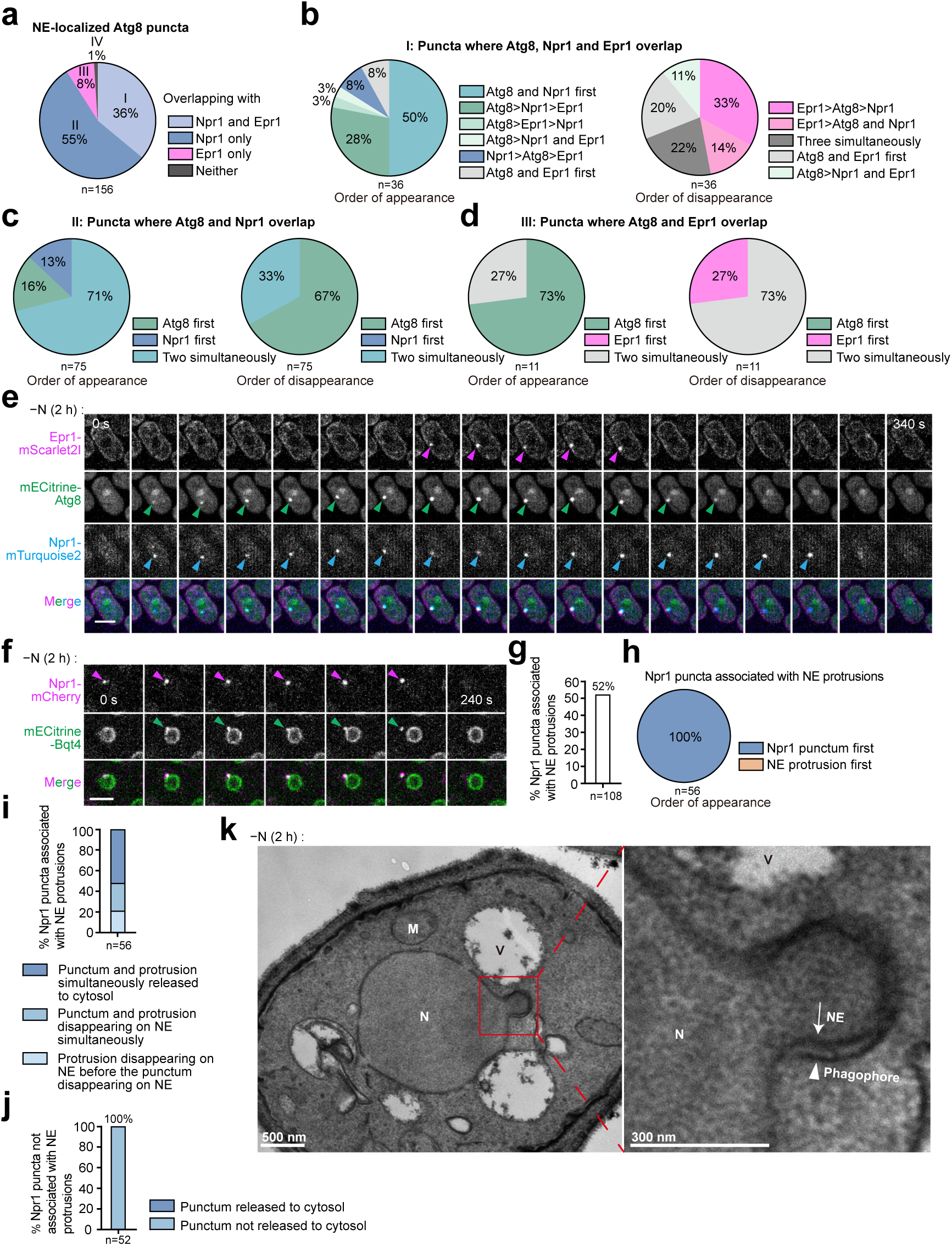
Dynamics of nucleophagy-related Atg8, Epr1, and Npr1 puncta and NE protrusions. (a) The overlap of NE-localized Atg8 puncta with Epr1 and Npr1 puncta in time-lapse imaging data. Time-lapse imaging was performed on cells co-expressing Epr1-mScarlet2I, mECitrine-Atg8, and Npr1-mTurquoise2 after 2 h of nitrogen starvation (20-s intervals). NE-localized Atg8 puncta, whose entire lifespans from appearance to disappearance were captured, were analyzed and categorized as: Type I (Epr1+/Npr1+), Type II (Npr1+ only), Type III (Epr1+ only), or Type IV (neither). (b-d) The order of appearance and disappearance of Npr1, Epr1, and Atg8 at types I (b), II (c), and III (d) NE-localized Atg8 puncta. (e) Representative time-lapse series of a cell containing a type I punctum. Epr1-mScarlet2I, mECitrine-Atg8 and Npr1-mTurquoise2 signals enriched at this punctum are denoted by arrowheads. Bar, 3 µm. (f) NE protrusion release into the cytosol. A representative time-lapse series showing that a Bqt4-labeled NE protrusion associated with an Npr1 punctum was released into the cytosol. Bar, 3 µm. (g) Quantification of the percentages of Npr1 puncta associated with NE protrusions in wild-type cells after 2 h of nitrogen starvation. A total of 108 Npr1 puncta, whose entire lifespans from appearance to disappearance were captured by time-lapse imaging, were analyzed. (h) The order of appearance of Npr1 puncta and associated NE protrusions. (i) Quantification of the percentages of Npr1 puncta and associated NE protrusions that were released into the cytosol or that disappeared from the NE without being detected in the cytosol. (j) None of the Npr1 puncta that were not associated with NE protrusions were released into the cytosol. (k) A representative electron microscopy image of starved wild-type cells showing a phagophore (arrowhead) wrapping around an NE protrusion. N: nucleus; V: vacuole; M: mitochondrion.

We analyzed the sequential appearance and disappearance of Npr1, Epr1, and Atg8 at types I-III NE-localized Atg8 puncta **(Fig. 4b-d)**. For type I puncta, Atg8 and Npr1 appeared simultaneously and before Epr1 in 50% of cases, while Atg8 appeared before both receptors in 34%. Regarding disappearance, Epr1 disappeared first in 47% of cases, while puncta of all three proteins disappeared simultaneously in 22%. Among type II puncta, Atg8 and Npr1 appeared simultaneously in 71% of cases, with Atg8 disappearing first in 67%. For type III puncta, Atg8 appeared first in 73% of cases, with Atg8 and Epr1 disappearing simultaneously in 73%. These results indicate that the sequential orders of the appearance and disappearance of Atg8, Npr1, and Epr1 puncta are not fixed.

All analyzed NE-localized Npr1 and Epr1 puncta (72 and 52, respectively) exhibited overlap with Atg8, consistent with the absence of these puncta in *atg8Δ* cells **(Supplementary Fig. 4g)**. Overall, in 64% of cases, Atg8 appeared simultaneously with the receptors, while in 26%, Atg8 preceded receptor appearance. The rare cases (10%) where Npr1 preceded visible Atg8 puncta suggest that Npr1 puncta formation may require only low levels of Atg8. Notably, Npr1 puncta often persisted after Atg8 disappearance, indicating that their stability does not require continuous high levels of Atg8.

### Nucleophagy receptors promote NE protrusion formation

During Atg39-mediated nucleophagy in *S. cerevisiae*, the NE forms outward protrusions^13,16,31^. To investigate whether NE protrusions also form during nucleophagy in *S. pombe*, we imaged nitrogen-starved cells expressing Npr1-mCherry and the INM protein mECitrine-Bqt4 **(Fig. 4f)**. Time-lapse analysis revealed that in 52% of cases, Npr1 puncta formation was followed by a short outward projection of the mECitrine-Bqt4 signal at the same position, indicating NE protrusion formation **(Fig. 4f, g)**. Notably, we did not observe any instances where NE protrusions formed before the emergence of Npr1 puncta at the same sites **(Fig. 4h)**. Furthermore, the nucleoplasmic protein Pus1-mTurquoise2 co-localized with Bqt4 projections, indicating the presence of nucleoplasmic components within the NE protrusions **(Supplementary Fig. 4h)**.

Similar observations were made with Epr1, where time-lapse analysis of cells expressing Epr1-ymScarlet2I and mECitrine-Bqt4 showed that in 54% of cases, the formation of NE-localized Epr1 puncta was associated with the subsequent formation of NE protrusions at the same sites **(Supplementary Fig. 4i-k)**. These data suggest that the assembly of Npr1 and Epr1 puncta at the NE frequently precedes and potentially initiates the formation of NE protrusions.

Next, we investigated the relationship between Atg8 puncta and NE protrusions during nucleophagy. Time-lapse imaging of cells expressing ymScarlet2I-Atg8 and mECitrine-Bqt4 revealed that the formation of NE-localized Atg8 puncta was frequently followed by the formation of NE protrusions at the same locations **(Supplementary Fig. 4n-p)**. Electron microscopy analysis confirmed the presence of NE protrusions in nitrogen-starved cells **(Fig. 4k and Supplementary Fig. 4s)**, revealing that these NE protrusions were closely surrounded by membranes that terminated at the necks of the protrusions—likely representing phagophores where lipidated Atg8 resides.

To assess the role of nucleophagy receptors in NE protrusion formation, we compared the frequency of Atg8-positive NE protrusions in wild-type and *epr1Δ npr1Δ* cells. We found that 40% of NE-localized Atg8 puncta were associated with NE protrusions in wild-type cells, compared to only 4% in *epr1Δ npr1Δ* cells **(Supplementary Fig. 4o)**. These findings indicate that nucleophagy receptors play a critical role in promoting the formation of NE protrusions during nucleophagy.

### Distinct dynamics of nucleophagy components during NE protrusion release

To investigate the fate of NE protrusions formed during nucleophagy, we analyzed time-lapse imaging data, focusing on sites where Atg8 and nucleophagy receptors assembled into puncta **(Fig. 4f, i, j and Supplementary Fig. 4i, l-n, q, r)**. Using the INM protein mECitrine-Bqt4 as an NE marker, we observed that the disappearance of an NE protrusion was often accompanied by the emergence of a nearby Bqt4-labeled punctum in the cytosol **(Fig. 4f)**. This suggests that NE protrusions are released into the cytosol.

Next, we investigated the dynamics of Npr1, Epr1, and Atg8 puncta in relation to NE protrusion release. For Npr1 puncta associated with NE protrusions, 52% exhibited simultaneous release with the NE protrusion into the cytosol **(Fig. 4i)**. Of the remaining 48%, 27% disappeared simultaneously with the NE protrusion, and 21% disappeared after the protrusion had vanished, without subsequent detection of cytosolic puncta. The released Npr1 puncta likely represent autophagosomes that have fully engulfed nuclear cargo and detached from the NE during nucleophagy. Notably, Npr1 puncta not associated with NE protrusions were never observed to be released into the cytosol **(Fig. 4j)**, suggesting that NE protrusion formation is essential for successful nucleophagy-related autophagosome generation.

Epr1 puncta associated with NE protrusions exhibited three patterns: (1) simultaneous release with NE protrusions into the cytosol (30%); (2) disappearance before NE protrusion release (25%); and (3) disappearance of both from the NE without cytosolic puncta detection (45%, including 36% simultaneous disappearance and 9% NE protrusion disappearance first) **(Supplementary Fig. 4l)**. Similar to Npr1, Epr1 puncta without associated NE protrusions were never released into the cytosol **(Supplementary Fig. 4m)**.

The behavior of Atg8 puncta associated with NE protrusions differed notably—none were released simultaneously with NE protrusions into the cytosol. In 53% of instances, Atg8 puncta disappeared from the NE before the release of NE protrusions. In the remaining 47%, both disappeared from the NE, either simultaneously (20%) or with Atg8 puncta disappearing first (27%) **(Supplementary Fig. 4q)**. Atg8 puncta not associated with NE protrusions were never observed to be released into the cytosol **(Supplementary Fig. 4r)**.

Taken together, these findings reveal distinct behaviors among nucleophagy components during NE protrusion release. Atg8 typically disappears from the NE prior to protrusion release, while Npr1 tends to be released concurrently. Epr1 shows an intermediate pattern, being released together with NE protrusions in about half of the protrusion release events. Since NE protrusion release likely follows NE fission and autophagosome closure, these findings suggest that Atg8 is usually recycled before autophagosome closure, likely through Atg4-mediated delipidation.^32^ Conversely, Npr1, being an integral membrane protein, always remains within autophagosomes. As a peripheral membrane protein, Epr1 may escape from autophagosomes before their closure. Importantly, we observed two types of premature termination of nucleophagy in wild-type cells: the presence of nucleophagy receptor puncta without associated NE protrusions and the disappearance of NE protrusions without detectable cytosolic puncta, indicating that nucleophagy can abort naturally at either NE protrusion formation or release.

### Npr1- and Epr1-mediated nucleophagy maintains nuclear morphology and survival during nitrogen starvation

It is known that *S. cerevisiae* mutants defective in nucleophagy exhibit abnormal nuclear morphology during nitrogen starvation^12^, and that *S. pombe* autophagy mutants display aberrant nuclear morphology during meiosis, which is induced by nitrogen starvation^33^. To investigate the role of nucleophagy in maintaining normal nuclear morphology during nitrogen starvation in *S. pombe*, we examined NE morphology using the INM protein mECitrine-Bqt4 as a marker.

Under nutrient-rich conditions, NE morphology appeared normal in both *atg5Δ* and *epr1Δ npr1Δ* mutants **(Fig. 5a and Supplementary Fig. 5a)**. However, after 24 hours of nitrogen starvation, these mutants exhibited striking NE aberrations characterized by projections with intensified mECitrine-Bqt4 signals. These abnormal NE structures fell into two categories: extended NE projections, where the projections extend away from the NE, and ring-shaped NE projections, where both ends of a projection are associated with the NE. In the *atg5Δ* mutant, extended NE projections were more frequently observed than ring-shaped NE projections, whereas the *epr1Δ npr1Δ* mutant predominantly displayed ring-shaped projections. Neither type of aberration was observed in nitrogen-starved wild-type, *epr1Δ*, or *npr1Δ* cells **(Fig. 5a and Supplementary Fig. 5a)**. This suggests that nucleophagy redundantly promoted by Npr1 and Epr1 is required for maintaining normal nuclear morphology during nitrogen starvation. Supporting this, re-introducing either Npr1 or Epr1, but not their AIM-mutated forms, into the *epr1Δ npr1Δ* mutant restored normal NE morphology **(Fig. 5a and Supplementary Fig. 5a)**.

**Fig. 5:**
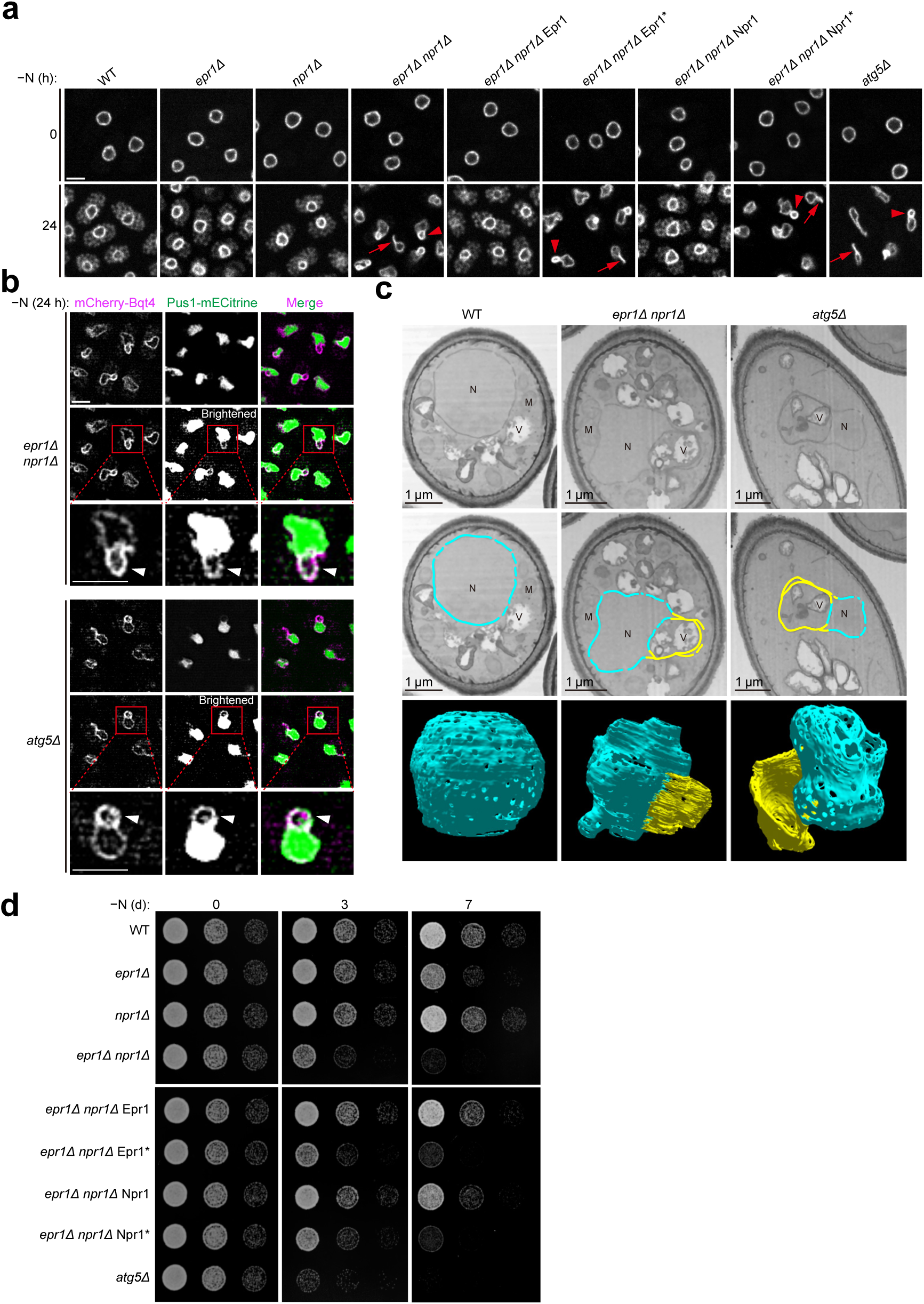
Npr1- and Epr1-mediated nucleophagy is essential for maintaining nuclear morphology and cell survival under nitrogen starvation. (a) *epr1Δ npr1Δ and atg5Δ* cells formed NE projections during nitrogen starvation. This phenotype in *epr1Δ npr1Δ* was rescued by reintroducing either Epr1 or Npr1 in an AIM-dependent manner. Fluorescence microscopy was used to visualize cells expressing the INM protein mECitrine-Bqt4 before and after 24 h of nitrogen starvation. Representative ring-shaped NE projections are indicated by arrowheads, and extended NE projections by arrows. Bar, 3 µm. (b) The nucleoplasmic protein Pus1-mECitrine colocalized with mCherry-Bqt4 at NE projections in *epr1Δ npr1Δ* and *atg5Δ* cells. Images were processed by deconvolution. Weak Pus1-mECitrine signals at NE projections became clearly visible only after brightness adjustments. In magnified views of the boxed areas, NE projections are indicated by arrowheads. Bar, 3 µm. (c) FIB-SEM analysis revealed the three-dimensional structures of NE projections in *epr1Δ npr1Δ* and *atg5Δ* cells. Top: representative FIB-SEM slices of a wild-type (WT) cell, an *epr1Δ npr1Δ* cell, and an *atg5Δ* cell. Middle: the same slices with the NE colored blue and NE projections colored yellow. N, nucleus; M, mitochondrion; V, vacuole. Bottom: 3D reconstructions of the NE and NE projections in the cells shown in the top and middle rows. Approximately 300 FIB-SEM slices were used for each reconstruction. The NE is colored blue, and the NE projections are colored yellow. (d) Reduced survival of *epr1Δ npr1Δ* and *atg5Δ* cells after nitrogen starvation. This phenotype of *epr1Δ npr1Δ* was alleviated by reintroducing either Epr1 or Npr1 in an AIM-dependent manner. Cells subjected to nitrogen starvation for 0, 3, and 7 days were plated in five-fold serial dilutions on YES plates, which were photographed after colony formation.

To further characterize these NE projections, we examined the localization of the nucleoplasmic protein Pus1-mECitrine after 24 hours of nitrogen starvation **(Fig. 5b)**. Pus1-mECitrine colocalized with mCherry-Bqt4 at the NE projections in both *atg5Δ* and *epr1Δ npr1Δ* cells, indicating that these projections are nucleoplasm-containing structures **(Fig. 5b)**. The intensified Bqt4 signals observed at these projections likely result from the presence of two layers of NE sandwiching a thin layer of nucleoplasm.

To visualize the ultrastructures of the NE projections, we employed focused ion beam-scanning electron microscopy (FIB-SEM) **(Fig. 5c)**. Wild-type cells exhibited a spherical NE shape **(Fig. 5c and Supplementary Video 1)**. In stark contrast, and consistent with our light microscopy observations, *epr1Δ npr1Δ* and *atg5Δ* cells exhibited aberrant, non-spherical NE morphology featuring prominent projections. FIB-SEM slices through these projections confirmed the presence of two layers of NE in the projections **(Fig. 5c)**. Additionally, three-dimensional reconstruction revealed that the ring-shaped NE projections observed under light microscopy correspond to cross-sections of dome-like or semi-dome-like structures extending from the NE surface **(Fig. 5c and Supplementary Video 2, 3)**.

Next, we investigated the contribution of nucleophagy to cell survival during nitrogen starvation using spot assays **(Fig. 5e)**. After 3 days of nitrogen starvation, both *epr1Δ npr1Δ* and *atg5Δ* mutants showed reduced survival compared to wild-type cells, while *epr1Δ* or *npr1Δ* mutants were unaffected. This survival defect in *epr1Δ npr1Δ* cells was rescued by re-introducing either Npr1 or Epr1, but not their AIM-mutated forms.

Collectively, these results demonstrate that Npr1- and Epr1-mediated nucleophagy plays essential roles in maintaining normal nuclear morphology and promoting cell survival during nitrogen starvation in *S. pombe*.

### Inhibition of nucleophagy by Lem2 overexpression

During our examination of various INM proteins as NE markers, we made a serendipitous discovery that the INM protein Lem2, when expressed exogenously from a medium-strength promoter (*P41nmt1*), nearly completely blocked the autophagic processing of Npr1-mECitrine and Pus1-mECitrine **(Fig. 6a)**. Notably, this inhibition was specific to cargos of Npr1- and Epr1-mediated nucleophagy, as no effect was observed on the autophagic processing of the nucleoporin mECitrine-Nup82 **(Supplementary Fig. 6a)** and the cortical ER membrane protein Rtn1-mECitrine **(Supplementary Fig. 6b)**. These findings suggest that elevated Lem2 expression selectively disrupts nucleophagy.

**Fig. 6:**
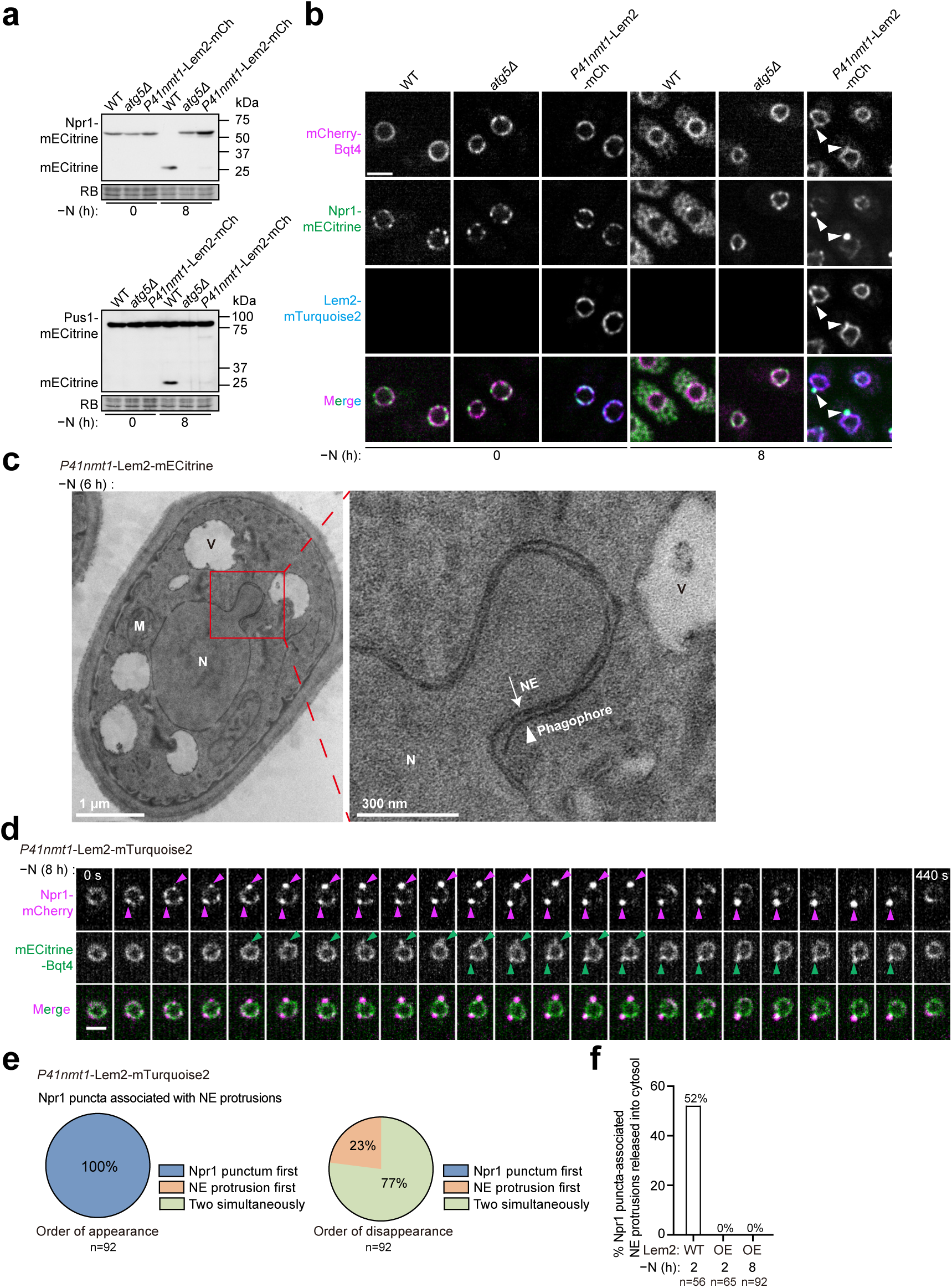
Lem2 overexpression inhibits nucleophagy. (a) Overexpression of Lem2-mCherry (mCh) from the exogenous *P41nmt1* promoter inhibited nucleophagy. Autophagic processing of Npr1-mECitrine and Pus1-mECitrine was analyzed by immunoblotting in wild-type (WT), *atg5Δ*, and Lem2-overexpressing cells. (b) During nitrogen starvation, Npr1 puncta associated with NE protrusions (arrowheads) formed in Lem2-overexpressing cells. Bar, 3 µm. (c) A representative electron microscopy image of Lem2-overexpressing cells after 6 h of nitrogen starvation shows a phagophore (arrowhead) wrapping around an NE protrusion. N, nucleus; V, vacuole; M, mitochondrion. (d) Time-lapse fluorescence microscopy showed that Npr1 puncta (magenta arrowheads) in Lem2-overexpressing cells exhibited kinetics of appearance and disappearance similar to those in wild-type cells. mECitrine-Bqt4 served as an NE marker, and NE protrusions are indicated by green arrowheads. Fluorescence images were captured at 20-second intervals. Bar, 2 µm. (e) The order of appearance and disappearance of Npr1 puncta and associated NE protrusions was analyzed in Lem2-overexpressing cells after 8 h of nitrogen starvation. A total of 92 Npr1 puncta associated with NE protrusions were examined. (f) Quantification of the percentages of Npr1 puncta-associated NE protrusions released into the cytosol during time-lapse analysis. A total of 56 Npr1 puncta-associated NE protrusions in WT cells after 2 h of nitrogen starvation, 65 in Lem2-overexpressing (OE) cells after 2 h of nitrogen starvation, and 92 in Lem2-overexpressing (OE) cells after 8 h of nitrogen starvation were analyzed.

To determine the level of Lem2 required to inhibit nucleophagy, we expressed Lem2-mCherry using four different promoters with increasing expression strengths: *P81nmt1* < *Padf1* < *P41nmt1* < *Pcyc1* **(Supplementary Fig. 6c, d)**. Interestingly, even a modest level of exogenous Lem2 expression (2-4-fold of the endogenous level) driven by the weakest promoter, *P81nmt1*, resulted in observable, albeit moderate, inhibition of nucleophagy. Expression from stronger promoters (*Padf1* and above), at levels 8-16-fold or higher than the endogenous level, resulted in near-complete inhibition of nucleophagy. These findings indicate that Lem2 inhibits nucleophagy in a dose-dependent manner.

To explore how this inhibition occurs, we examined the localization of the nucleophagy receptor Npr1 in Lem2-overexpressing cells. After 8 hours of nitrogen starvation, wild-type cells showed Npr1-mECitrine primarily in the vacuole lumen, while both *atg5Δ* and Lem2-overexpressing cells retained Npr1-mECitrine at the NE **(Fig. 6b)**. Notably, Lem2-overexpressing cells displayed NE-localized Npr1-mECitrine puncta, unlike *atg5Δ* cells where no such puncta were observed. This indicates that Lem2 overexpression permits the initial phase of nucleophagy—Npr1 puncta formation—but interferes with later stages. Similar to wild-type cells, Npr1-mECitrine puncta in Lem2-overexpressing cells were often associated with NE protrusions **(Fig. 6b)**, which also contained the nucleoplasmic protein Pus1-mECitrine **(Supplementary Fig. 6e)**. Electron microscopy revealed that NE protrusions in Lem2-overexpressing cells were surrounded by phagophores **(Fig. 6c and Supplementary Fig. 6f)**, indicating they are nucleophagy intermediate structures.

Time-lapse imaging showed that in both wild-type and Lem2-overexpressing cells, NE-localized Npr1 puncta were dynamic structures with lifetimes of several minutes **(Fig. 6d and Supplementary Fig. 6h)**. These puncta were often associated with NE protrusions that formed after the puncta appeared (**Fig. 6d, e and Supplementary Fig. 6g**). However, while more than half of the NE protrusions in wild-type cells eventually detached from the NE and were released into the cytosol **(Fig. 4i, 6f)**, NE protrusions in Lem2-overexpressing cells never detached. Instead, they disappeared from the NE without being released into the cytosol **(Fig. 6d, f)**. In most cases, NE protrusions disappeared simultaneously with Npr1 puncta, although in 23% of instances, NE protrusions disappeared before Npr1 puncta **(Fig. 6e)**. These findings indicate that Lem2 overexpression allows nucleophagy to progress to the stage of NE protrusion formation but blocks the subsequent step of protrusion release.

To further explore this phenomenon, we analyzed the behavior of Atg8 in Lem2-overexpressing cells. While Atg8 and Npr1 co-localized at NE puncta, appearing simultaneously in most cases **(Supplementary Fig. 6h-j)**, Atg8 puncta consistently disappeared from the NE before Npr1 puncta **(Supplementary Fig. 6j)**. Furthermore, Atg8 puncta always disappeared before the NE protrusions **(Supplementary Fig. 6k)**, suggesting that components of the autophagy machinery dissociate before the retraction of NE protrusions during the abortive nucleophagy process.

### Chromatin-INM tethering inhibits nucleophagy

To investigate how Lem2 overexpression inhibits nucleophagy, we analyzed which regions of Lem2 are critical for this effect. Lem2 contains an N-terminal LEM domain (amino acids 1-60) that binds DNA^34^, a Bqt4-binding motif (BBM, amino acids 261-279) that localizes Lem2 to the NE by interacting with the INM protein Bqt4^35^, and two transmembrane helices **(Fig. 7a)**.

**Fig. 7:**
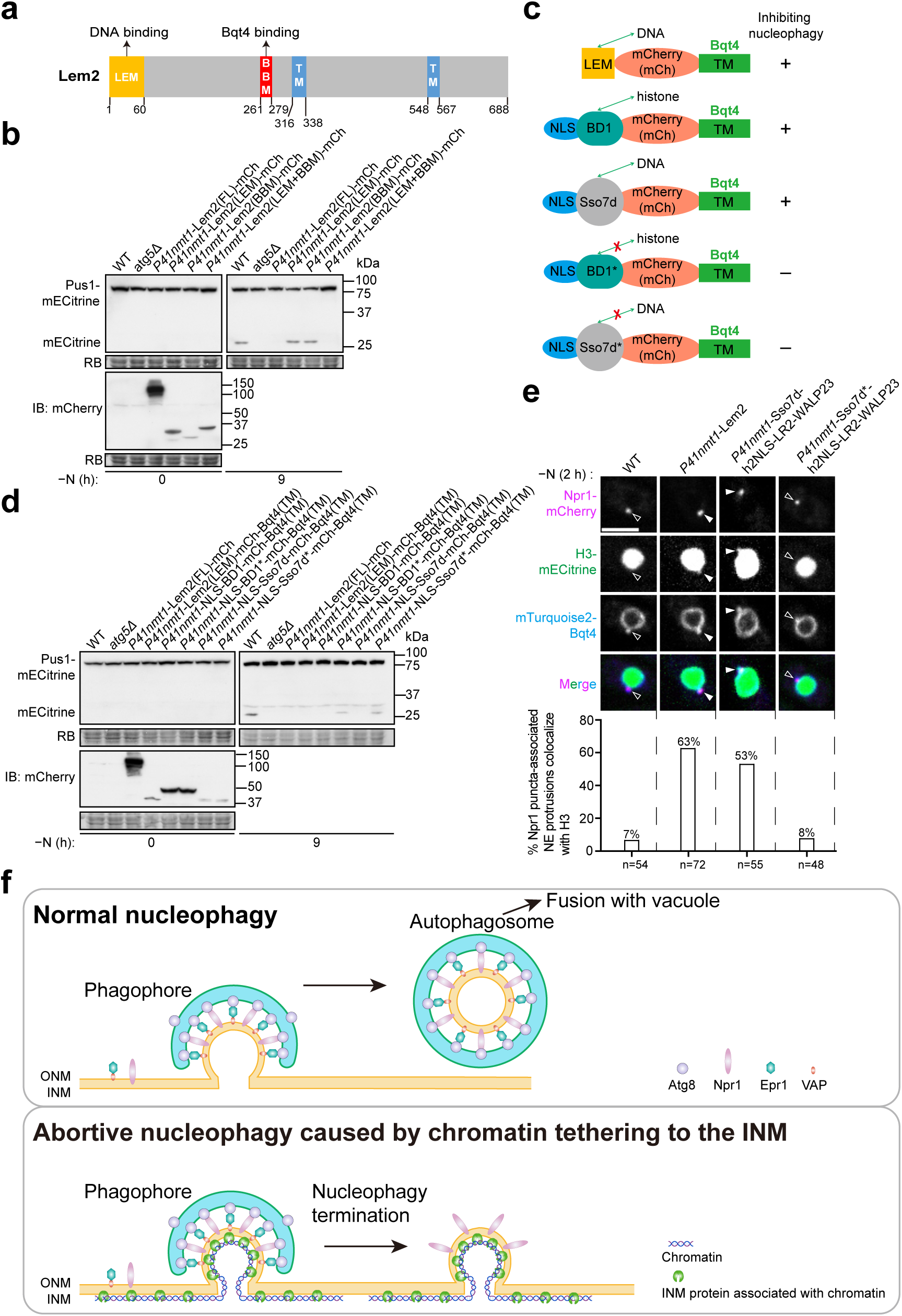
Chromatin tethering to the INM inhibits nucleophagy. (a) Schematic of the functional domains and transmembrane helices (TMs) of Lem2. Two functional domains, the LEM domain (LEM) and the Bqt4-binding motif (BBM), are highlighted. (b) A fusion of the LEM and BBM domains of Lem2, but not either domain alone, inhibits nucleophagy. FL, full length; mCh, mCherry. (c) Schematic of fusion proteins containing a DNA- or histone-binding domain from Lem2, Bdf1, or Sso7d, fused to the C-terminal transmembrane helix (TM) of Bqt4 (amino acids 412-432). Each fusion protein also includes a nuclear localization signal (NLS) and mCherry. BD1 refers to the first histone-binding bromodomain of Bdf1 (amino acids 66-208), while BD1* carries a Y123F mutation that disrupts histone binding. Sso7d is a nonspecific DNA-binding protein from *Archaea*, while Sso7d* carries W24A and R43E mutations that impair its DNA-binding ability. (d) Fusion proteins containing the DNA- or histone-binding domains of Lem2, Bdf1, or Sso7d fused to the TM of Bqt4 inhibit nucleophagy in a DNA/histone binding-dependent manner. The expression levels of these mCherry (mCh)-tagged fusion proteins were analyzed by immunoblotting using an anti-mCherry antibody. (e) Histone H3-mECitrine frequently co-localized with Npr1 puncta-associated NE protrusions in Lem2-overexpressing cells and Sso7d-h2NLS-LR2-WALP23-expressing cells. In contrast, wild-type cells and Sso7d*-h2NLS-LR2-WALP23-expressing cells exhibited minimal co-localization. h2NLS-LR2-WALP23 is an artificial INM protein comprising the NLS from the budding yeast protein Heh2 (h2NLS), a random sequence linker (LR2), and an artificially designed transmembrane helix peptide WALP23. Top: representative images in which Npr1 puncta-associated NE protrusions with co-localized H3 signals are indicated by solid arrowheads, while those without co-localized H3 signals are indicated by hollow arrowheads. Bar, 3 µm. Bottom: quantification of the percentages of Npr1 puncta-associated NE protrusions co-localized with H3. (f) Schematic models illustrating normal nucleophagy (top) and abortive nucleophagy resulting from chromatin tethering to the INM (bottom).

Using truncation analysis, we found that an N-terminal soluble fragment of Lem2 (amino acids 1-279), which includes the LEM domain and BBM, was sufficient to inhibit nucleophagy, whereas fragments lacking either domain were ineffective **(Supplementary Fig. 7a)**. Internal deletion analysis confirmed that both domains are necessary for nucleophagy inhibition **(Supplementary Fig. 7a)**. Furthermore, a fusion construct containing just the LEM domain and BBM inhibited nucleophagy, while neither domain alone was sufficient **(Fig. 7b)**. These results establish that the combination of the LEM domain and BBM represents the minimal functional unit required for nucleophagy inhibition.

Because the LEM domain binds DNA and the BBM binds Bqt4, we hypothesized that their combination tethers chromatin to the INM, where Bqt4 resides. To test this hypothesis, we substituted the BBM in the fusion construct with the C-terminal transmembrane helix (TM, amino acids 412-432) of Bqt4, a single-pass INM protein with its N-terminus facing the nucleoplasm^36^. This LEM-Bqt4(TM) fusion localized to the NE **(Supplementary Fig. 7d)** and effectively inhibited nucleophagy **(Fig. 7c, d)**, suggesting that the role of the BBM in nucleophagy inhibition is to mediate INM association.

To determine whether the LEM domain’s role is chromatin association, we tested whether it could be replaced by other chromatin-binding domains. We substituted the LEM domain with either the histone-binding bromodomain (BD1) of *S. pombe* Bdf1 or the non-specific DNA-binding protein Sso7d from *Archaea* **(Fig. 7c, d)**^37,38^. Fusion proteins containing BD1 or Sso7d, along with the TM of Bqt4, localized to the NE **(Supplementary Fig. 7d)** and effectively inhibited nucleophagy **(Fig. 7d)**. Notably, when the histone-binding ability of BD1 was disrupted by a point mutation (Y123F, BD1*)^39^, or the DNA-binding ability of Sso7d was weakened by two mutations (W24A and R43E, Sso7d*)^40^, these fusion proteins no longer strongly inhibited nucleophagy **(Fig. 7d)**. These results indicate that the main role of the LEM domain in nucleophagy inhibition is to mediate chromatin association.

To further solidify the role of chromatin-INM tethering, we tested Sso7d fused to the multi-transmembrane INM protein Bqt3, a binding partner of Bqt4^36^. This fusion also effectively inhibited nucleophagy **(Supplementary Fig. 7b, d)**. Notably, even though the known physiological functions of Bqt3 and Bqt4 require their interaction with each other^36,41^, nucleophagy inhibition by Sso7d-Bqt3 occurred in the absence of *bqt4* and nucleophagy inhibition by Sso7d-Bqt4(TM) occurred in the absence of *bqt3* **(Supplementary Fig. 7b-d)**, suggesting that nucleophagy inhibition does not depend on Bqt3-Bqt4 interaction. Remarkably, the Sso7d-Bqt3 fusion protein was able to inhibit nucleophagy even when expressed from much weaker promoters, including the *P81nmt1* promoter **(Supplementary Fig. 7e)**.

To rule out the possibility that nucleophagy inhibition depends on specific functions of Bqt4(TM) or Bqt3 beyond their INM localization, we employed an artificial INM protein h2NLS-LR2-WALP23, consisting of the NLS of budding yeast Heh2, a random linker sequence LR2, and an artificial transmembrane helix WALP23^42^. Fusion proteins combining BD1 or Sso7d with h2NLS-LR2-WALP23 inhibited nucleophagy, whereas fusion proteins with the mutated forms BD1* or Sso7d* did not, despite all constructs localizing to the NE **(Supplementary Fig. 7f, g)**. This definitively demonstrates that simply tethering chromatin to the INM is sufficient to inhibit nucleophagy.

A key feature of nucleophagy inhibition by Lem2 overexpression is the blockage of NE protrusion release into the cytosol **(Fig. 6h)**. We hypothesized that this blockage is caused by excessive chromatin tethering to the INM, leading to the presence of chromatin in the NE protrusions. Analysis of histone H3 localization revealed its presence in 7% of Npr1 puncta-associated NE protrusions in wild-type cells, increasing to 63% in cells overexpressing Lem2 **(Fig. 7e)**. Similarly, H3 was frequently detected in Npr1 puncta-associated NE protrusions in cells expressing Sso7d-h2NLS-LR2-WALP23 but not in cells expressing Sso7d*-h2NLS-LR2-WALP23 **(Fig. 7e)**. Super-resolution microscopy confirmed H3 localization within Npr1-positive structures in Lem2-overexpressing cells **(Supplementary Fig. 7h)**. These findings suggest that chromatin presence in NE protrusions prevents their release, resulting in abortive nucleophagy.

## Discussion

In this study, we identified Npr1, an Atg8-binding protein that localizes to the ONM and acts as a dedicated nucleophagy receptor. During nitrogen starvation, Npr1 functions redundantly with Epr1 to mediate the selective degradation of nuclear components. Both Npr1 and Epr1 assemble into Atg8-positive NE-localized puncta where NE protrusions containing nucleoplasmic material form. While many of these protrusions are eventually released into the cytosol, presumably as cargo enclosed in autophagosomes, some do not undergo this release. These abortive nucleophagy events may be caused by the presence of chromatin within the protrusions, as artificially tethering chromatin to the INM strongly inhibits nucleophagy by preventing the release of NE protrusions **(Fig. 7f)**.

All currently known autophagy receptors involved in nucleophagy exhibit relatively narrow species distributions. Atg39 is exclusively found in budding yeast species within the family *Saccharomycetaceae*, but not in the sister family *Saccharomycodaceae*, indicating that it originated no earlier than 152 million years ago^43^. Similarly, Epr1 is confined to the fission yeast genus *Schizosaccharomyces*^18^, whose origin is dated around 207 million years ago^43^. Even more restricted is Npr1, which is found only in *S. pombe* and is absent in other fission yeast species (annotated by PomBase as an *S. pombe* specific protein)^44^, suggesting that its emergence occurred no earlier than 108 million years ago^43^. This frequent emergence of nucleophagy receptors during evolution may be explained by their minimal functional requirements— specifically, the presence of an AIM and localization to the ONM—as evidenced by the successful functional replacement of Epr1 and Npr1 with artificial fusion proteins that localizes to the ONM and contains a cytosol-facing AIM.

The co-existence of Epr1 and Npr1 in *S. pombe*, despite their redundant roles in nucleophagy during nitrogen starvation, may be explained by their non-overlapping functions in other cellular contexts. Notably, Epr1 is essential for ER stress-induced ER-phagy and nucleophagy^18,19^. It is conceivable that Npr1 plays a critical role in nucleophagy under specific circumstances that were not examined in this study, highlighting the need for further investigation into its functions.

In *S. cerevisiae*, Atg39’s role in nucleophagy extends beyond merely acting as a nucleophagy receptor. Through its single transmembrane helix, which spans the ONM, and its C-terminal amphipathic helices, which are located in the NE lumen and bind to the INM, Atg39 establishes a physical connection between the ONM and the INM, facilitating NE deformation during nucleophagy^13,16^. In contrast, Npr1 and Epr1 appear to function solely as autophagy receptors, as their roles in nucleophagy can be substituted by either of two ONM-localized membrane proteins (Kms1 and Erg11) artificially fused with an AIM. In *S. pombe*, the NE-deforming function attributed to Atg39 in *S. cerevisiae* is likely carried out by other, yet-to-be-identified protein(s) involved in nucleophagy.

Previous studies have established that Atg39 is dispensable for the autophagic degradation of nuclear pore components in *S. cerevisiae*, with Nup159 instead acting as an autophagy receptor to facilitate this process^14-16^. Consistent with this, our study reveals that the autophagic degradation of nuclear pore components in *S. pombe* is independent of Epr1 and Npr1. Notably, our TurboID-Atg8 proximity labeling experiments showed enrichment of Nup146 **(Supplementary Table 1)**, the *S. pombe* ortholog of *S. cerevisiae* Nup159. Further studies are needed to determine whether Nup146 serves as an autophagy receptor in *S. pombe*.

In Atg39-mediated nucleophagy in *S. cerevisiae*, two different models have been proposed for how NE fission gives rise to nucleus-derived vesicles that are enveloped within autophagosomes. One model posits that the INM and ONM protrude together towards the cytosol at the site where Atg39 assembles, subsequently undergoing simultaneous fission^13^. The alternative model suggests a two-step NE fission process: first, the INM undergoes fission, leading to the formation of INM-derived vesicles within the NE lumen; second, the ONM undergoes fission, a process that is partially dependent on the fission factor Dnm1^16,31^. Our EM analysis revealed that in *S. pombe*, the INM and ONM protrude together, suggesting that the simultaneous fission model may be applicable to *S. pombe*. The fission of the ONM may involves Yep1, the ortholog of human REEP1-4 proteins and *S. cerevisiae* Atg40^19^. However, the identity of the INM fission factor(s) and the mechanism coupling ONM and INM fission remain unclear.

In *S. cerevisiae*, nucleophagy defects resulting from *atg39* deletion impair cell viability during nitrogen starvation^12^. This phenotype has been attributed to compromised Atg39-mediated degradation of Nvj1, a key micronucleophagy factor whose accumulation triggers excessive micronucleophagy^45^. In this study, we found that in *S. pombe*, nucleophagy defects caused by the deletion of *epr1* and *npr1* similarly result in decreased cell viability under nitrogen starvation. However, since *S. pombe* lacks Nvj1 and micronucleophagy has not been observed in this organism, the mechanisms driving viability loss are likely distinct. Notably, in *epr1Δ npr1Δ* mutant cells, we observed aberrant nuclear morphology characterized by membrane structures extending from the NE surface. The cause of this phenotype and the relationship between this phenotype and cell viability loss remain unclear. We speculate that an imbalance in the supply and turnover of NE components may underlie the observed NE extensions.

Our finding that the presence of chromatin in NE protrusions leads to abortive nucleophagy reveals how cells safeguard genomic integrity during the degradation of nuclear components. While this underscores a protective mechanism, the precise molecular basis for chromatin-mediated termination of nucleophagy remains unclear. The failure of protrusions to release into the cytosol suggests that NE fission at the protrusion neck is inhibited. This inhibition could arise from two potential mechanisms: chromatin may passively obstruct membrane remodeling at the fission site, or dedicated surveillance systems may actively detect chromatin within protrusions and suppress NE fission. Further studies are needed to distinguish between these passive and active regulatory models.

## Methods

### Strain and plasmid construction

The *S. pombe* strains used in this study are listed in Supplementary Table 2, and the plasmids used in this study are listed in Supplementary Table 3. Unless stated otherwise, the strains were cultured in EMM medium at 30 °C. The compositions of the EMM medium, EMM−N medium, and other media are as decribed^46^. The strains were constructed as previously described^46^. Deletion strains were generated through PCR-based gene targeting. The strains with Npr1 or Lem2 endogenously tagged at their C-termini were constructed using PCR-based tagging^47^.

Plasmids expressing proteins fused with TurboID under the *P41nmt1* promoter were constructed using modified pDUAL vectors^20,48^. The other protein-expressing plasmids were based on stable integration vectors (SIVs)^49,50^. These plasmids allow integration at specific loci, namely *ura4*, *ade6*, *lys3*, or *his5*. The plasmid expressing histone H3 (Hht2)-yeGFP was derived from the plasmid pDB5568^51^, which was based on the pAde6^PmeI^ SIV plasmid and used a unique PmeI site as the linearization site. Other plasmids, designed to express proteins fused with various N-terminal or C-terminal tags (mCherry, mScarlet2I, ymScarlet2I, mECitrine, or mTurquoise2), were based on modified SIV plasmids with a unique NotI site as the linearization site^50^. Plasmids expressing Npr1/Npr1(W22A/V25A)-mECitrine under its endogenous promoter were constructed by introducing the *npr1* promoter, together with the coding sequence of Npr1/Npr1(W22A/V25A), into a modified SIV containing the sequence encoding mECitrine. Plasmids expressing AIM^art^-Npr1(30-244)-mCherry, AIM^art^-Man1-mCherry, Rtn1-AIM^art^-mCherry, AIM^art^-mCherry-Kms1, AIM^art^-Erg11-mCherry, and Erg11-AIM^art^-mCherry were constructed by inserting the coding sequences of Npr1(30-244), Man1, Rtn1, Kms1, and Erg11, respectively, into a modified SIV containing the *P41nmt1* promoter and the sequence encoding AIM^art^-mCherry. Plasmids expressing mCherry-Kms1-AIM^art^ were constructed by inserting the coding sequence of Kms1-AIM^art^ into a modified SIV containing the *P41nmt1* promoter and the sequence encoding mCherry. AIM^art^ corresponds to 3xEEEWEEL^29,30^. The plasmid expressing Lem2(LEM+BBM)-mECitrine was constructed by inserting the coding sequences of amino acids 1-60 and amino acids 261-279 of Lem2 into a modified SIV containing the *P41nmt1* promoter and the sequence encoding mECitrine^34,35^. Plasmids used to investigate the roles of LEM and BBM, including Lem2(LEM)-mCherry-Bqt4(TM), NLS-BD1-mCherry-Bqt4(TM), NLS-BD1*-mCherry-Bqt4(TM), NLS-Sso7d-mCherry-Bqt4(TM), NLS-Sso7d*-mCherry-Bqt4(TM), NLS-BD1-mCherry-h2NLS-LR2-WALP23, NLS-BD1*-mCherry-h2NLS-LR2-WALP23, NLS-Sso7d-mCherry-h2NLS-LR2-WALP23, and NLS-Sso7d*-mCherry-h2NLS-LR2-WALP23, were constructed by inserting the coding sequences of Bqt4(412-432) and h2NLS-LR2-WALP23 into modified SIVs, each containing the *P41nmt1* promoter and the sequences encoding Lem2(1-60), NLS-BD1, NLS-BD1(Y123F), NLS-Sso7d, or NLS-Sso7d(W24A/R43E), and mCherry^36-42^. The nucleotide sequence encoding NLS is ATGCCTAAGAAGAAGCGTAAGGTC. Plasmids that encode NLS-Sso7d-mCherry-Bqt3 under the endogenous promoter of Bqt3 were generated by inserting the *bqt3* promoter and the coding sequence of Bqt3 into a modified SIV containing the NLS-Sso7d and mCherry sequences. Plasmids expressing NLS-Sso7d-mCherry-Bqt3 under exogenous promoters were constructed by inserting the *P81nmt1* promoter, *P41nmt1* promoter, or *Pcyc1* promoter, along with the coding sequence of Bqt3, into a modified SIV containing the NLS-Sso7d and mCherry sequences.

Modified SIVs were used in this study to express proteins fused with GFP_1-10_ and 7×GFP_11_^23-25^. The plasmid expressing Sum3-GFP_1-10_-mCherry under the *Padh1* promoter was constructed by inserting the coding sequences of Sum3 and GFP_1-10_ into a modified SIV containing the *Padh1* promoter and mCherry sequences. The plasmid expressing Gbs1-GFP_1-10_-mCherry under the *Padh1* promoter was constructed by inserting the sequences of GFP_1-10_ and mCherry between codons 496 and 497 of Gbs1 and placing the coding sequence of the fusion protein downstream of the *Padh1* promoter in an SIV plasmid. Plasmids expressing 7×GFP_11_-Erg11, Erg11-7×GFP_11_, 7×GFP_11_-Npr1, and Npr1-7×GFP_11_ under the *P41nmt1* promoter were constructed by inserting the coding sequences of Erg11 or Npr1, together with the sequence encoding 7×GFP_11_, into a modified SIV containing the *P41nmt1* promoter.

### TurboID-based proximity labeling and mass spectrometry analysis

Approximately 1000 OD_600_ units of TurboID-mCherry and TurboID-mCherry-Atg8 cells were collected following 4 hours of nitrogen starvation. The collected cells were washed three times with deionized water and subsequently centrifuged to remove the supernatant. The resulting pellet was then reconstituted in 20 mL of deionized water, followed by the addition of 20 mL of 0.7 M sodium hydroxide. The mixture was incubated at room temperature on a rolling wheel for 10 minutes. The alkaline-treated cells were centrifuged to remove the supernatant. Afterward, 1 mL of lysis buffer (2% SDS, 0.06 M Tris-HCl, 5% glycerol, and 4% 2-mercaptoethanol, pH 6.8) was added to resuspend the pellet, which was then incubated at 42°C for 20 minutes. The supernatant was collected after centrifugation at 16,246 × g for 30 minutes. For the purification of biotinylated proteins, approximately 100 µL of Streptavidin Agarose Resin (Thermo Fisher Scientific, Cat#20359) was used. The streptavidin beads were pre-washed twice using washing buffer A (50 mM Tris-HCl, 150 mM NaCl, 1 mM EDTA, 1 mM EGTA, 1% Triton X-100, 0.4% SDS, 1% NP40, and 1× Roche protease inhibitor cocktail, pH 7.5). After the addition of the streptavidin beads, the supernatant was incubated at room temperature on a rolling wheel for 3 hours to allow the biotinylated proteins to bind to the streptavidin beads.

After incubation, the streptavidin beads were pelleted and washed twice with 1 mL of washing buffer A for 5 minutes and then incubated with 1 mL of washing buffer B (50 mM Tris-HCl, 2% SDS, pH 7.5) for 10 minutes. Subsequently, the beads were incubated twice with 1 mL of washing buffer A for 5 minutes each. Following this, biotinylated proteins were eluted from the beads by incubating the beads twice with 200 µL of elution buffer (50 mM Tris-HCl, 2% SDS, 5 mM biotin, pH 8.0) at 60°C for 20 minutes, using a ThermoMixer C (Eppendorf) set at 1,000 × rpm.

To precipitate the proteins, 100 µL of 100% trichloroacetic acid (TCA) was added to approximately 400 µL of eluate. The mixture was then incubated overnight at 4°C. Subsequently, the precipitated proteins were centrifuged at 16,246 × g at 4°C for 30 minutes, and the pellet was washed with acetone three times. The pellet was then resuspended with 30 µL of dissolution buffer (8 M urea, 100 mM Tris-HCl, pH 8.5) and dissolved by sonication using a water bath sonicator at room temperature for 15 minutes. Next, the proteins were reduced using 5 mM tris(2-carboxyethyl)phosphine at room temperature for 20 minutes, followed by alkylation with 10 mM iodoacetamide at room temperature for 15 minutes. The sample was diluted by a factor of 4 and digested into peptide fragments using trypsin at 37°C overnight. To terminate the trypsin digestion, formic acid was added to a final concentration of 5%.

After the completion of the digestion process, LC-MS/MS analysis was conducted using an Easy-nLC II HPLC instrument (Thermo Fisher Scientific), which was coupled to a Q Exactive Orbitrap mass spectrometer (Thermo Fisher Scientific). A total of 8 µL of peptides were loaded onto a pre-column (100 µm ID, 4 cm long, packed with C18 10 µm 120 Å resin from YMC Co., Ltd) and separated on an analytical column (75 µm ID, 10 cm long, packed with Luna C18 1.8 µm 100 Å resin from Welch Materials) using an acetonitrile gradient from 0% to 30% over a duration of 100 minutes. The flow rate during the separation was maintained at 250 nL/min. From each full scan (resolution 70,000), the top 15 most intense precursor ions were selected for higher-energy collisional dissociation tandem mass spectrometry (HCD MS2) analysis, with a normalized collision energy of 27 and a dynamic exclusion time of 30 seconds. The fragment ions obtained from the tandem mass spectrometry were detected using the Orbitrap in normal scan mode. Charge state rejection was enabled, and unassigned charge states as well as charge states 1, 7, 8, and >8 were rejected. Mass spectrometry data were analyzed using pFind software, with a peptide false discovery rate (FDR) cutoff of 1%^52^.

### Immunoprecipitation

For immunoprecipitation (IP), approximately 100 OD_600_ units of log-phase cells were collected and subjected to three washes with water. The cell pellet was mixed with 100 µL of lysis buffer A (50 mM HEPES, 1 mM EDTA, 150 mM NaCl, 10% glycerol, 3 mM DTT, 3 mM PMSF, Roche 3×Protease inhibitor cocktail, pH 7.5) and 800 µL of glass beads (BioSpec) with a diameter of 0.5 mm for cell lysis. The cells were lysed using the FastPrep-24 instrument at a speed of 6.5 m/sec for 20 seconds. This lysis process was repeated three times. Then, 300 µL of lysis buffer B (50 mM HEPES, 1 mM EDTA, 150 mM NaCl, 10% glycerol, 0.05% NP40, 1 mM DTT, 1 mM PMSF, Roche 1×Protease inhibitor cocktail, pH 7.5) was added to the cell lysate. After centrifugation, 20 µL of the supernatant was retained as input, while the remainder was added to pre-washed GFP-Trap agarose beads at 4°C for 3 hours. The beads had been pre-washed with lysis buffer B. After centrifugation, the agarose beads were washed twice with wash buffer (50 mM HEPES, 1 mM EDTA, 150 mM NaCl, 10% glycerol, 0.05% NP40, 1 mM DTT, pH 7.5) and twice with lysis buffer B. The proteins bound to the GFP-Trap agarose beads were eluted by incubating them in SDS loading buffer (60 mM Tris-HCl, 4% SDS, 4% 2-mercaptoethanol, 5% glycerol, 0.002% bromophenol blue, pH 6.8) at 42°C for 20 minutes for subsequent immunoblotting analysis.

### Fluorescence microscopy

We used cells cultured in liquid medium (EMM or EMM−N) for microscopy analysis. Live-cell imaging was performed using a Dragonfly 201-40 high-speed spinning-disk confocal microscope (Andor Technology), equipped with a 100×/1.4 NA objective lens, a Sona sCMOS camera, and two filter sets for mCherry/YFP/CFP and mCherry/GFP, respectively. Super-resolution images were acquired using a High Intelligent and Sensitive Structured Illumination Microscope (HIS-SIM, CSR Biotech Co., Ltd, Guangzhou), equipped with a 100×/1.5 NA objective lens, an sCMOS Flash 4.0 V2 camera, and a filter set for mCherry/GFP. SIM images were collected and analyzed as described^53^. The microscopy images obtained were analyzed using FIJI^54^.

For time-lapse imaging, a suspension of nitrogen-starved cells cultured in liquid medium (EMM−N) was applied to an agar pad placed on a microscope glass slide (7.5 cm long)^55^. Firstly, two double-sided tapes were placed in the middle third of a clean glass slide, about 1.5 cm apart. Then, approximately 50 μL of hot melted agar (in EMM−N) was placed onto the glass slide, and immediately a coverslip (2.2 cm long) was placed on top of the agar drop. After the agar pad solidified in 3-5 minutes, the coverslip was removed, and 1 μL of the cell suspension was placed onto the agar pad. Time-lapse imaging analysis was performed after placing a new coverslip on top of the agar pad and ensuring the coverslip adhered tightly to the glass slide through the double-sided tapes.

### Immunoblotting-based protein processing assay

A total of 5 OD_600_ units of cells were collected for lysis. The cells were resuspended in 300 µL of 20% TCA, and 700 µL of glass beads with a diameter of 0.5 mm were added. The cells were then lysed using the FastPrep-24 instrument at a speed of 6.5 m/sec for 20 seconds, repeating this process three times. The cell lysates were transferred into new centrifuge tubes through centrifugation. To adjust the TCA concentration to 10%, 300 µL of deionized water was added to the cell lysates. The mixture was vortexed and centrifuged at 865 × g for 30 minutes. The resulting pellet was resuspended in SDS loading buffer (60 mM Tris-HCl, 4% SDS, 4% 2-mercaptoethanol, 5% glycerol, 0.002% bromophenol blue, pH 6.8) and incubated at 42°C for 20 minutes. Subsequently, centrifugation was performed at 16,246 × g for 5 minutes, and the resulting supernatant was separated by 10% SDS-PAGE and subjected to immunoblotting using specific antibodies. The antibodies used for immunoblotting were anti-GFP mouse monoclonal antibody (1:3,000 dilution, Roche, Cat#11814460001) and anti-mCherry rabbit polyclonal antibody (1:3,000 dilution, Invitrogen, Cat#PA5-34974). Post-immunoblotting staining of the PVDF membrane using Reactive Brown 10 (RB) served as the loading control^56^.

### Electron microscopy

For conventional transmission electron microscopy (TEM) analysis, a total of 50 OD_600_ units of cells were collected after either 2 or 6 hours of nitrogen starvation. The cells were fixed with glutaraldehyde and KMnO_4_^57^. Following fixation, the cells underwent 13 rounds of water washing to remove any brownish particles. They were then dehydrated by passing through a series of graded ethanol solutions. Ultimately, the dehydrated samples were embedded in Spurr’s resin^57^. Thin sections of 90 nm were examined using an FEI Tecnai G2 Spirit electron microscope equipped with a Gatan 895 4k × 4k CCD camera.

For electron microscopy analysis utilizing the genetically encoded EM tag MTn, samples containing 20 OD_600_ units of vegetative cells expressing Npr1-mECitrine-MTn were processed following the previously described method^18,26^.

For focused ion beam-scanning electron microscopy (FIB-SEM), a total of 50 OD_600_ units of cells were collected after 24 hours of nitrogen starvation. Cell samples were processed using the same method as that used for preparing conventional TEM samples. The resin-embedded samples were mounted on aluminum stubs. FIB-SEM datasets were acquired using a Zeiss Crossbeam 550 microscope equipped with ATLAS 3D software (ZEISS). During the milling process with the focused gallium-ion beam, a milling current of 700 pA at 30 kV was used from the gallium emitter. The resin-embedded samples were milled in 10 nm layers. Scanning EM images were captured using an SE2 detector set at 2 kV and 1 nA. The image resolution in the xy plane was 10 nm/pixel. The alignment of image stacks, visualization, 3D reconstructions, and creation of movies were carried out using the Dragonfly Pro software (version 2022.2).

### Yeast two-hybrid (Y2H) assay

The yeast two-hybrid (Y2H) analysis was conducted using the Matchmaker system 3 (Clontech) to express two fusion proteins, namely the bait and prey. Bait plasmids were constructed by inserting the coding sequence of Atg8 into the pGBKT7 vector. Prey plasmids were constructed by inserting the coding sequence of Npr1/Npr1(W22A/V25A) into the pGADT7 vector. The AH109 yeast strain was co-transformed with the bait and prey plasmids and then selected on double dropout medium (SD/−Leu/−Trp). The activation of the *HIS3* and *ADE2* reporter genes was evaluated using quadruple dropout medium (SD/−Leu/−Trp/−His/−Ade). Photographs were taken after incubating the transformants on double dropout medium and quadruple dropout medium at 30°C for 3 to 4 days.

### Growth phenotype assay (spot assay)

The *S. pombe* strains for the spot assay were inoculated into EMM medium supplemented with histidine, leucine, and uracil. The cells were cultured at 30°C until they reached the logarithmic phase. A portion of the cells was collected, while the remaining cells were subjected to nitrogen starvation for 3 days and 7 days, respectively. Subsequently, the harvested cells were subjected to a five-fold dilution and spotted onto YES plates. The plates were then incubated at 30°C and photographed after 3 days.

### Prediction of transmembrane topology and protein complex structure

The transmembrane topology was predicted using the CCTOP web server (https://cctop.ttk.hu)^22^. The structure of the Atg8-Npr1 complex was predicted using AlphaFold2-Multimer (version 2.3.1) with default parameters^27,28^. The structure exhibiting the highest confidence score among the predicted outputs was chosen for subsequent analysis. The visual representation of the predicted structure was generated using the Mol* Viewer (version 4.2.0)^58^.

## Supplemental information

**Supplementary Table 1**. Excel file containing mass spectrometry results of the TurboID experiment.

**Supplementary Table 2**. Excel file listing *S. pombe* strains used in this study.

**Supplementary Table 3**. Excel file listing plasmids used in this study.

**Supplementary Video 1**. 3D FIB-SEM reconstruction of the NE in a wild-type cell.

The NE is shown in blue.

**Supplementary Video 2**. 3D FIB-SEM reconstruction of the NE, NE projection, and vacuoles within the NE projection in an *epr1Δ npr1Δ* cell.

The NE is shown in blue. The nuclear projection is shown in yellow. The vacuoles are shown in purple.

**Supplementary Video 3**. 3D FIB-SEM reconstruction of the NE, NE projection, and vacuoles within the NE projection in an *atg5Δ* cell.

The NE is shown in blue. The nuclear projection is shown in yellow. The vacuoles are shown in purple.

## Acknowledgements

We are grateful to Ying Liu and He-Xia Luo of the NIBS Electron Microscopy Facility for their assistance with EM analysis. We also thank Wen-Yi Huang, Yin-Hua Lin, and Ke Du of Guangzhou Computational Super-Resolution Biotech Co., Ltd. for their support in live-cell imaging using their commercial super-resolution microscope (HIS-SIM). Additionally, we thank Yi-Feng Jiang and Zhen-Hua Zhang of the ZEISS Microscopy Customer Center in Beijing for their help with focused ion beam-scanning electron microscopy (FIB-SEM) analysis. We extend our gratitude to Meng-Li Shi for her assistance in drawing the models shown in Fig. 7f. This work was supported by grants from the Ministry of Science and Technology of the People’s Republic of China, the Beijing municipal government, and Tsinghua University.

## Author contributions

Conceptualization: Z.-H.M. and L.-L.D.; Methodology and investigation: Z.-H.M., Z.-Q.P., Z.-D.J., G.-C.S., Y.H., F.S., C.-X.Z., Y.-F.J., M.-Q.D., and L.-L.D.; Writing – original draft: Z.-H.M. and L.-L.D.; Writing – review and editing: Z.-H.M. and L.-L.D.; Funding acquisition: M.-Q.D. and L.-L.D.

## Materials & Correspondence

Correspondence and requests for materials should be addressed to L.-L.D.

---

**Supplementary Fig. 1:**
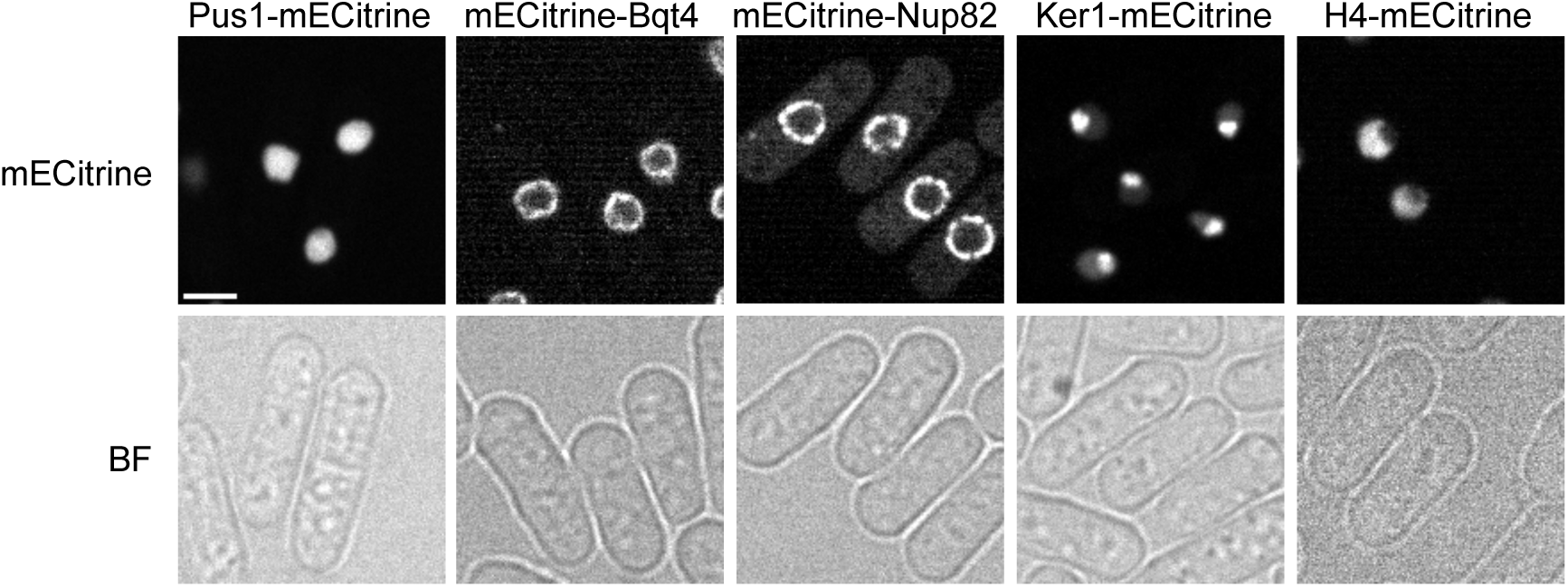
Subcellular localization of mECitrine-tagged nuclear proteins. Log-phase cells expressing mECitrine-tagged nuclear proteins, including Pus1-mECitrine, mECitrine-Bqt4, mECitrine-Nup82, Ker1-mECitrine, and H4-mECitrine, from the *P41nmt1* promoter were examined by fluorescence microscopy. BF, bright field. Bar, 3 µm.

**Supplementary Fig. 2:**
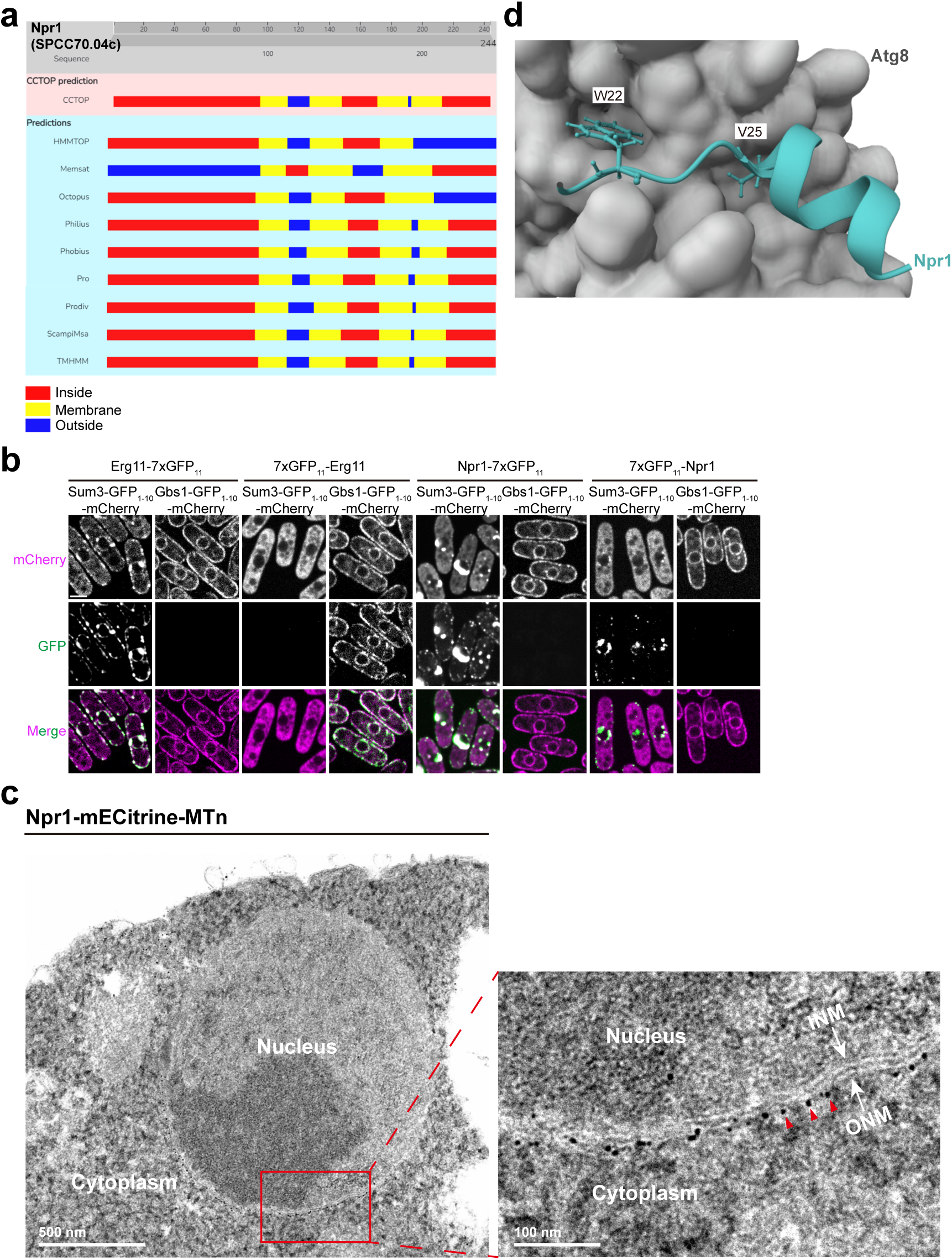
Identification of Npr1 as a candidate nucleophagy receptor. (a) The complete results of predicting the membrane topology of Npr1 (SPCC70.04c) using CCTOP. (b) Split GFP-based assays showed that the N- and C-termini of Npr1 are located in the cytosol. Log-phase cells were examined by fluorescence microscopy. GFP_1-10_ fused proteins were expressed from the *adh1* promoter, and GFP_11_ fused proteins were expressed from the *P41nmt1* promoter. Bar, 3 µm. (c) Electron microscopy (EM) analysis of gold nanoparticle-labeled Npr1. MTn tagging of Npr1 allowed labeling with EM-visible gold nanoparticles. A magnified view of the boxed area is shown on the right. INM: inner nuclear membrane; ONM: outer nuclear membrane. Arrowheads indicate three representative gold nanoparticles. (d) The AlphaFold2-Multimer-predicted structure of the protein complex between Npr1 and Atg8. Amino acids 21-32 of Npr1 are shown in a ribbon representation, with W22 and V25 in the AIM highlighted in a ball-and-stick representation. Atg8 is shown in a surface representation and colored grey.

**Supplementary Fig. 3:**
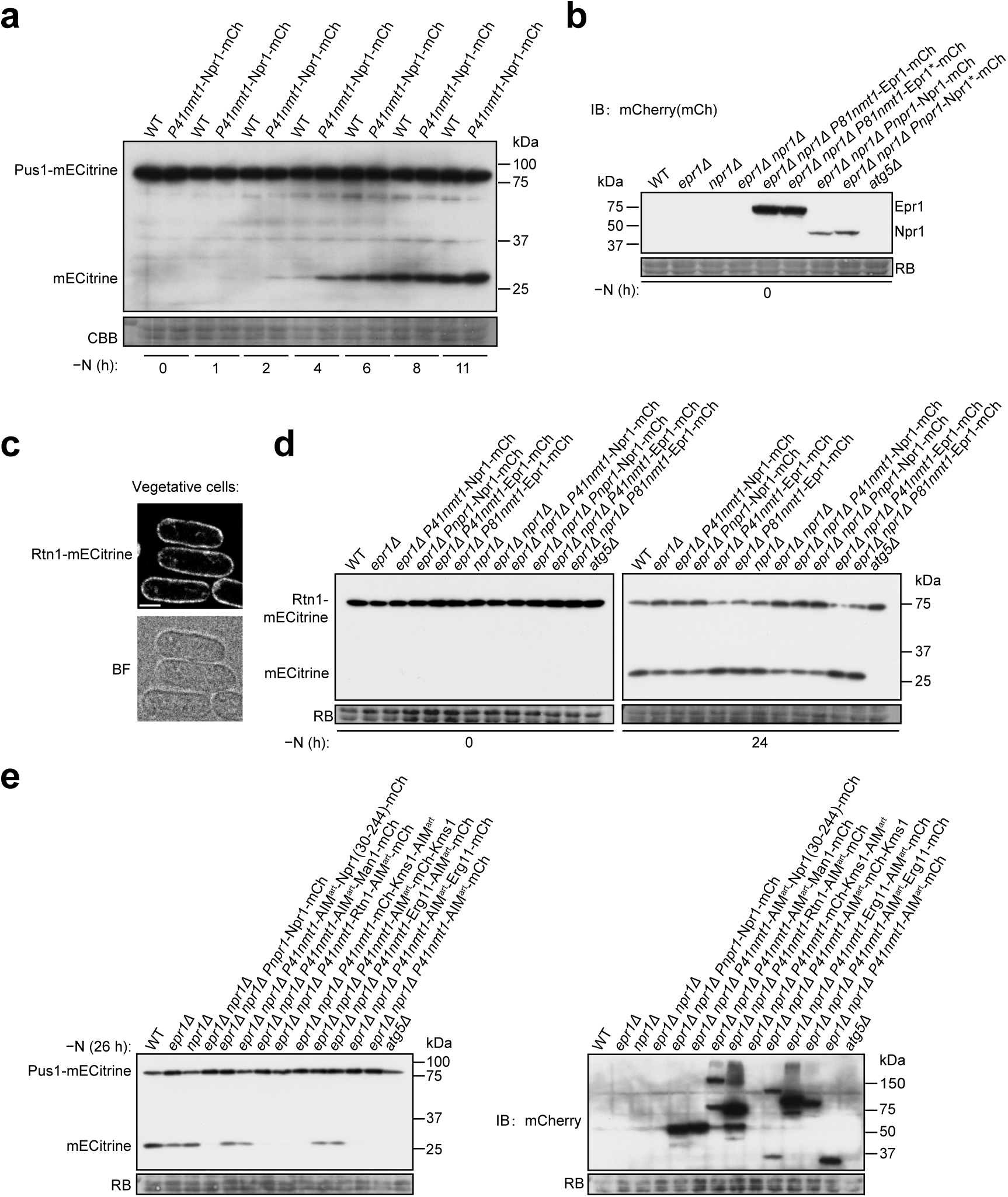
Npr1 and Epr1 are redundantly required for starvation-induced nucleophagy. (a) The overexpression of Npr1 through the addition of an extra copy of the *npr1* gene under the control of the *P41nmt1* promoter accelerated nucleophagy. (b) In the experiment shown in Fig. 3b, the expression levels of mCherry-tagged proteins expressed in *epr1Δ npr1Δ* were analyzed by immunoblotting using an anti-mCherry antibody. (c) Subcellular localization of the cortical ER membrane protein Rtn1-mECitrine. Log-phase cells expressing Rtn1-mECitrine from the *P41nmt1* promoter were examined by fluorescence microscopy. Bar, 3 µm. (d) The deletion of *epr1* resulted in a moderate defect in the autophagic degradation of the cortical ER membrane protein Rtn1 during nitrogen starvation, and this phenotype was not further exacerbated by the additional deletion of *npr1*. (e) Fusion of the artificial AIM (AIM^art^) to Man1, Rtn1, the lumen-facing C-terminus of Kms1, or the lumen-facing N-terminus of Erg11 did not rescue the nucleophagy defect of *epr1Δ npr1Δ* (left). The expression levels of the AIM-fused proteins tagged with mCherry were analyzed by immunoblotting using an anti-mCherry antibody (right).

**Supplementary Fig. 4:**
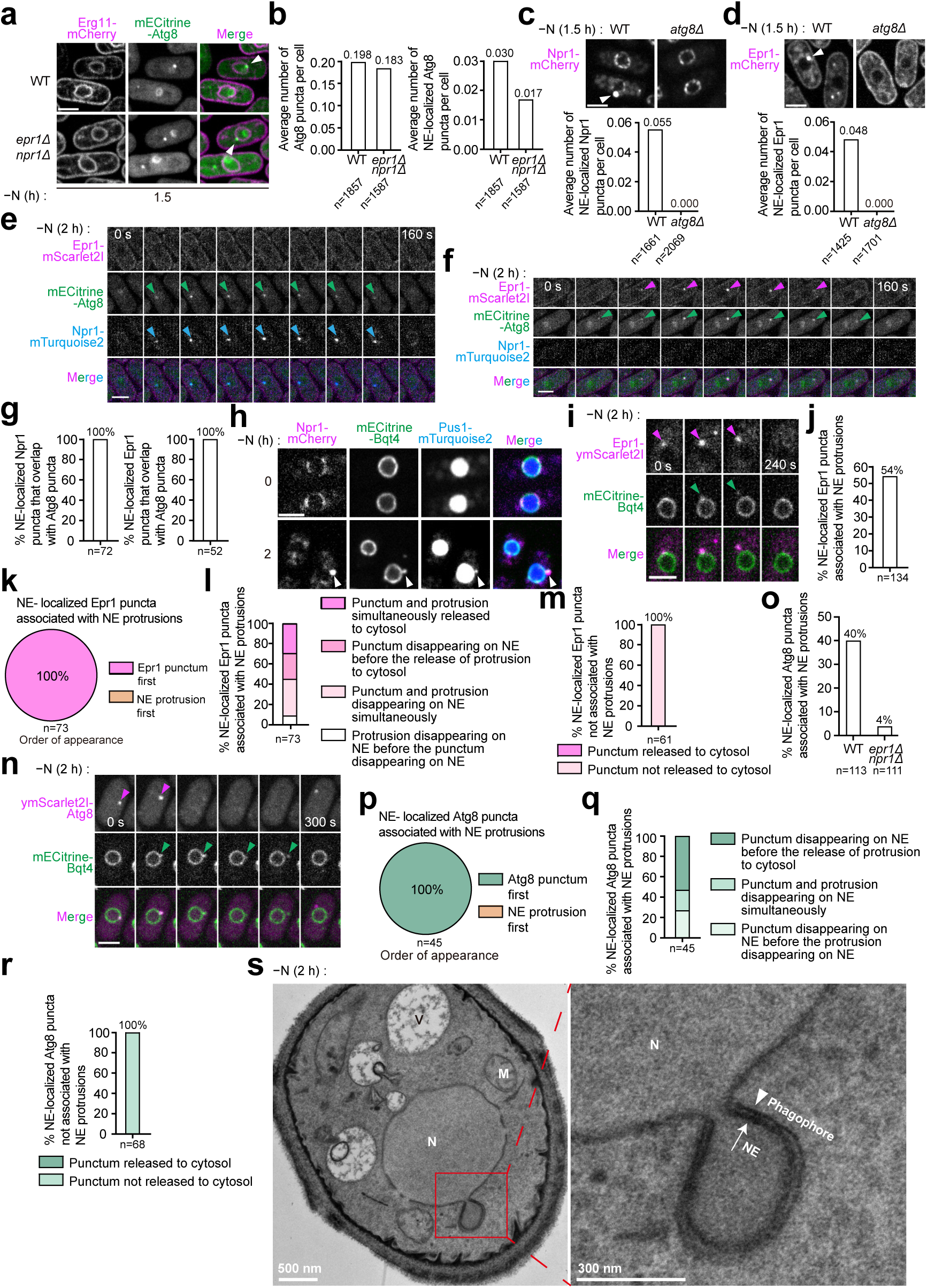
Epr1 and Npr1 puncta overlap with some but not all NE-localized Atg8 puncta, and NE protrusions are released into the cytosol. (a) NE-localized Atg8 puncta were observed in wild-type (WT) and *epr1Δ npr1Δ* cells after 1.5 h of nitrogen starvation. Erg11 was used as an ER membrane marker. NE-localized Atg8 puncta are indicated by white arrowheads. Bar, 3 µm. (b) Quantification of the average number of Atg8 puncta and NE-localized Atg8 puncta per cell in wild-type and *epr1Δ npr1Δ* cells from the experiment shown in (a). A total of 1857 wild-type cells and 1587 *epr1Δ npr1Δ* cells were analyzed. (c) NE-localized Npr1 puncta were not detected in *atg8Δ* cells after 1.5 h of nitrogen starvation. Top: representative images of cells containing NE-localized Npr1 puncta, indicated by white arrowheads. Bar, 3 µm. Bottom: quantification of the average number of NE-localized Npr1 puncta per cell. A total of 1661 wild-type cells and 2069 *atg8Δ* cells were analyzed. (d) NE-localized Epr1 puncta were not detected in *atg8Δ* cells after 1.5 h of nitrogen starvation. Top: representative images of cells containing NE-localized Epr1 puncta, indicated by white arrowheads. Bar, 3 µm. Bottom: quantification of the average number of NE-localized Epr1 puncta per cell. A total of 1425 wild-type cells and 1701 *atg8Δ* cells were analyzed. (e) Representative time-lapse images of a cell containing a type II punctum. Bar, 3 µm. (f) Representative time-lapse images of a cell containing a type III punctum. Bar, 3 µm. (g) Percentages of NE-localized Npr1 puncta (left) and Epr1 puncta (right) overlapping with Atg8 puncta in time-lapse imaging data. Cells co-expressing Epr1-mScarlet2I, mECitrine-Atg8, and Npr1-mTurquoise2 were subjected to fluorescence microscopy after 2 h of nitrogen starvation, with images acquired at 20-s intervals. A total of 72 NE-localized Npr1 puncta and 52 NE-localized Epr1 puncta were analyzed. Time-lapse imaging captured the entire lifespan of each punctum from appearance to disappearance. (h) The nucleoplasmic protein Pus1-mTurquoise2 localized at Npr1 puncta-associated NE protrusions in wild-type cells. Bar, 3 µm. (i) NE protrusion release into the cytosol. A representative time-lapse series showing that a Bqt4-labeled NE protrusion associated with an Epr1 punctum was released into the cytosol. Cells co-expressing Epr1-ymScarlet2I and mECitrine-Bqt4 were examined by fluorescence microscopy after 2 h of nitrogen starvation, with images acquired at 80-s intervals. Bar, 3 µm. (j) Quantification of the percentage of NE-localized Epr1 puncta associated with NE protrusions in wild-type cells after 2 h of nitrogen starvation. (k) The order of appearance of NE-localized Epr1 puncta and associated NE protrusions in wild-type cells after 2 h of nitrogen starvation. (l) Quantification of the percentages of NE-localized Epr1 puncta and associated NE protrusions released into the cytosol or disappearing from the NE in wild-type cells after 2 h of nitrogen starvation. (m) Quantification of the percentages of NE-localized Epr1 puncta not associated with NE protrusions that were released into the cytosol in wild-type cells after 2 h of nitrogen starvation. (n) An NE-localized Atg8 punctum disappeared from the NE before its associated NE protrusion was released into the cytosol. Cells co-expressing ymScarlet2I-Atg8 and the INM protein mECitrine-Bqt4 were examined by fluorescence microscopy after 2 h of nitrogen starvation, with images acquired at 60-s intervals. Bar, 3 µm. (o) Quantification of the percentages of NE-localized Atg8 puncta associated with NE protrusions in wild-type cells and *epr1Δ npr1Δ* cells after 2 h of nitrogen starvation. (p) The order of appearance of NE-localized Atg8 puncta and associated NE protrusions in wild-type cells after 2 h of nitrogen starvation. (q) Quantification of the percentages of NE-localized Atg8 puncta and associated NE protrusions released into the cytosol or simply disappearing from the NE in wild-type cells after 2 h of nitrogen starvation. (r) Quantification of the percentages of NE-localized Atg8 puncta not associated with NE protrusions that were released or not released into the cytosol in wild-type cells after 2 h of nitrogen starvation. (s) Electron microscopy analysis of wild-type cells treated with 2 h of nitrogen starvation showed a phagophore wrapping around an NE protrusion.

**Supplementary Fig. 5:**
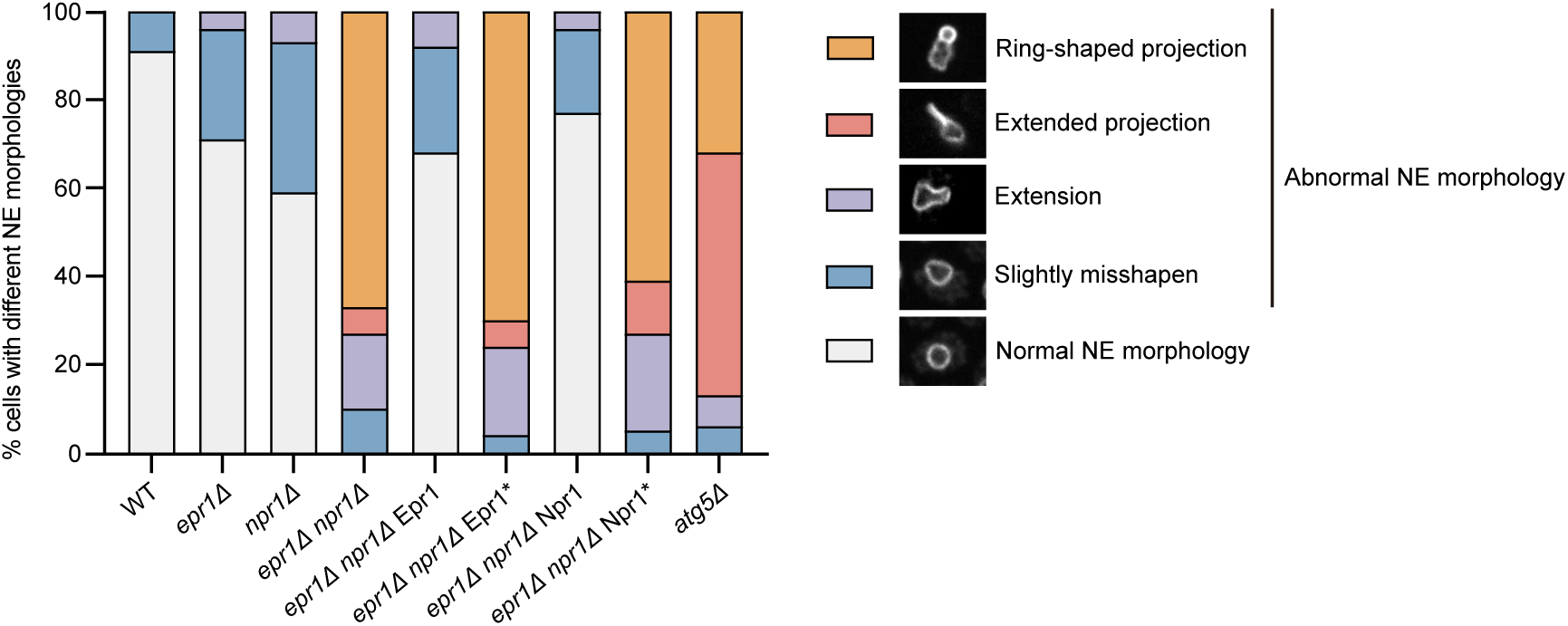
Npr1- and Epr1-mediated nucleophagy maintains nuclear morphology and survival during nitrogen starvation. Quantification of the percentages of cells exhibiting normal or various abnormal NE morphologies, as analyzed in Fig. 5a. Over 600 cells were examined per sample.

**Supplementary Fig. 6:**
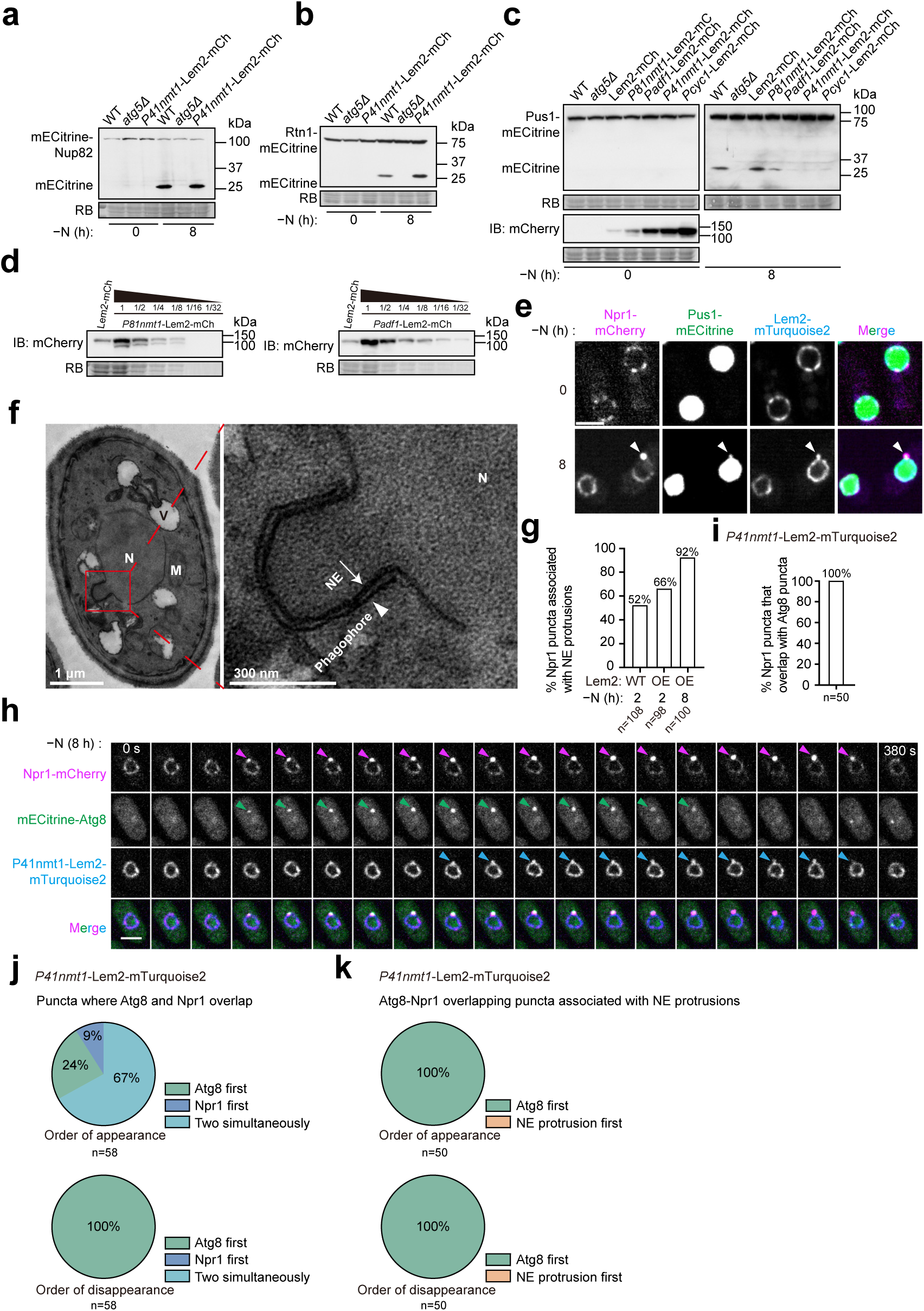
Inhibition of nucleophagy by Lem2 overexpression. (a) Overexpression of Lem2 does not inhibit the autophagic processing of mECitrine-Nup82. (b) Overexpression of Lem2 does not inhibit the autophagic processing of Rtn1-mECitrine. (c) The extent of nucleophagy inhibition dependents on the expression level of Lem2. Autophagic processing of Pus1-mECitrine was examined in cells with endogenously mCherry (mCh)-tagged Lem2 and in cells with exogenous Lem2-mCherry expressed from four different promoters using immunoblotting. The expression strengths of the promoters are as follows: *P81nmt1* < *Padf1* < *P41nmt1* < *Pcyc1.* Lem2-mCherry expression levels were analyzed by immunoblotting using an anti-mCherry antibody. (d) Lem2-mCherry expression levels under the control of *P81nmt1* and *Padf1* were 2-4 times and 8-16 times, respectively, that of the endogenously mCherry-tagged Lem2. Log-phase cells were used for the analysis. (e) In Lem2-mTurquoise2-overexpressing cells, the nucleoplasmic protein Pus1-mECitrine localized to Npr1 puncta-associated NE protrusions after 8 h of nitrogen starvation. Bar, 3 µm. (f) Electron microscopy analysis of Lem2-overexpressing cells treated with 6 h of nitrogen starvation showed a phagophore wrapping around an NE protrusion. (g) Quantification of the percentages of Npr1 puncta associated with NE protrusions. A total of 108 Npr1 puncta in wild-type (WT) cells treated with 2 h of nitrogen starvation, 98 Npr1 puncta in Lem2-overexpressing (OE) cells treated with 2 h of nitrogen starvation, and 100 Npr1 puncta in Lem2-overexpressing (OE) cells treated with 8 h of nitrogen starvation, were analyzed. Time-lapse imaging captured the entire lifespan of each punctum from appearance to disappearance. (h) Time-lapse analysis showed that in Lem2-mTurquoise2 overexpressing cells, Atg8 puncta colocalized with Npr1 puncta and the associated NE protrusions. Their appearance and disappearance followed kinetics similar to those of Atg8 puncta in wild-type cells. Cells co-expressing Npr1-mCherry, mECitrine-Atg8, and Lem2-mTurquoise2 were subjected to fluorescent microscopy after 8 h of nitrogen starvation, and fluorescence images were acquired at 20-s intervals. Bar, 3 µm. (i) The percentages of Npr1 puncta overlapping with Atg8 puncta were quantified from time-lapse imaging data. Cells co-expressing Npr1-mCherry, mECitrine-Atg8, and Lem2-mTurquoise2 were subjected to fluorescent microscopy after 8 h of nitrogen starvation, and fluorescence images were acquired at 20-s intervals. (j) The order of appearance and disappearance of Atg8 and Npr1 was analyzed at puncta where they overlapped in Lem2-mTurquoise2-overexpressing cells. (k) The order of appearance and disappearance of Atg8 and associated NE protrusions was analyzed at puncta where Atg8 and Npr1 overlapped in Lem2-mTurquoise2 overexpressing cells.

**Supplementary Fig. 7:**
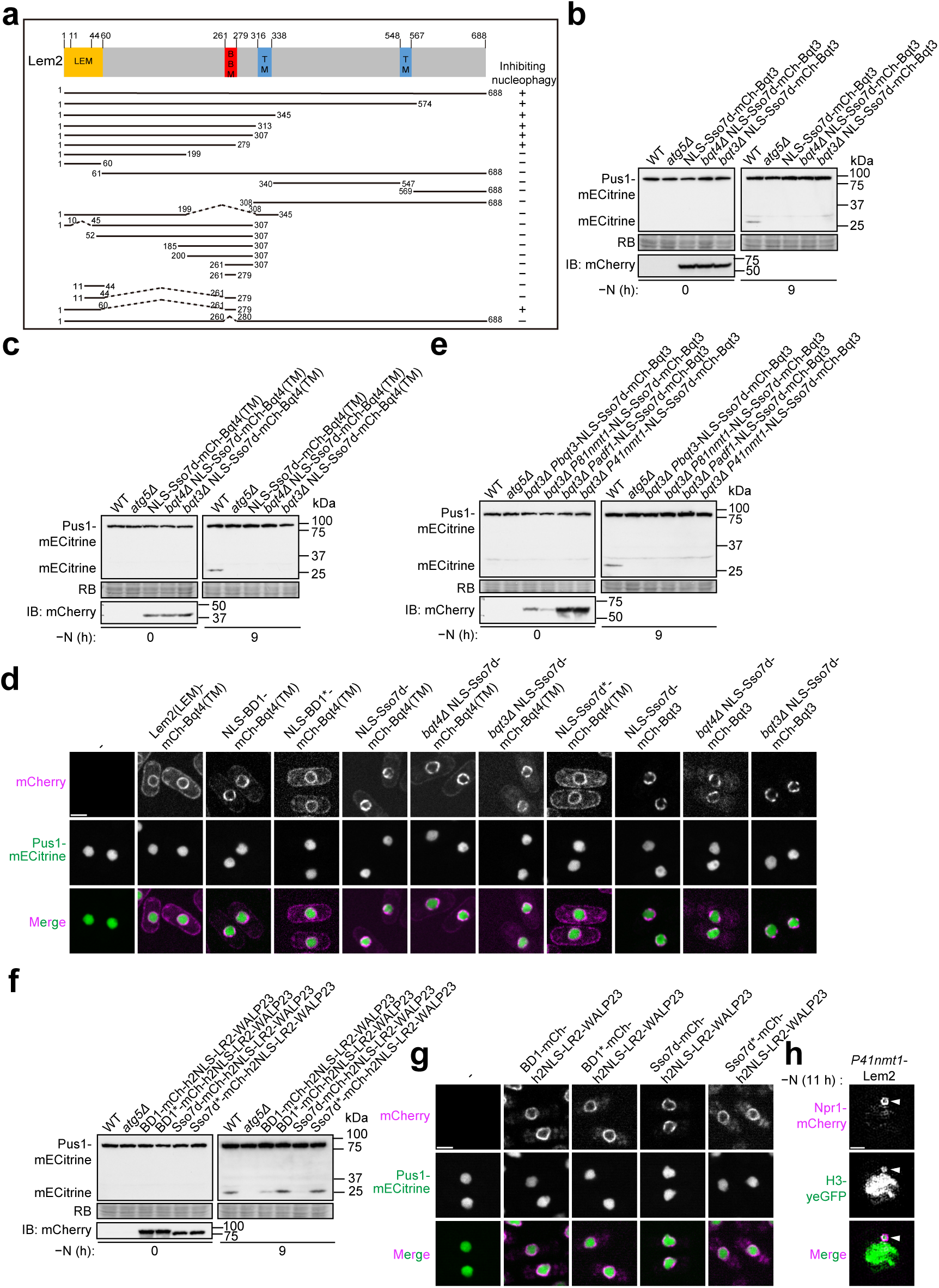
Chromatin-INM tethering inhibits nucleophagy. (a) Schematic showing the results of truncation and internal deletion analyses of Lem2, which identified the regions required for nucleophagy inhibition. Different forms of Lem2 with truncations or internal deletions were expressed under the *P41nmt1* promoter. Nucleophagy was assessed by monitoring the nitrogen starvation-induced relocalization of Npr1 to the vacuole. (b) The fusion protein between Sso7d and Bqt3, expressed from the *P41nmt1* promoter, inhibited nucleophagy even when the endogenous *bqt4* or *bqt3* gene was deleted. The fusion protein contains an NLS and mCherry (mCh). Expression levels were analyzed by immunoblotting with an anti-mCherry antibody. (c) The fusion protein between Sso7d and the C-terminal transmembrane helix (TM) of Bqt4, expressed from the *P41nmt1* promoter, also inhibited nucleophagy in the absence of the endogenous *bqt4* or *bqt3* gene. The fusion protein contains an NLS and mCherry (mCh). Expression levels were analyzed by immunoblotting with an anti-mCherry antibody. (d) Fusion proteins containing a DNA- or histone-binding domain from Lem2, Bdf1, or Sso7d fused to the C-terminal transmembrane helix (TM) of Bqt4 or full-length Bqt3 localized to the NE. Log-phase cells expressing a fusion protein and the nucleoplasmic protein Pus1-mECitrine were examined by fluorescence microscopy. Bar, 3 µm. (e) The fusion protein between Sso7d and Bqt3 inhibited nucleophagy even when expressed from the weak *P81nmt1* promoter in *bqt3Δ* cells. The fusion protein contains an NLS and mCherry (mCh). Expression levels were analyzed by immunoblotting with an anti-mCherry antibody. (f) Targeting BD1 or Sso7d to the INM using h2NLS-LR2-WALP23 inhibited nucleophagy. (g) Log-phase cells expressing a chromatin-tethering fusion protein and the nucleoplasmic protein Pus1-mECitrine were examined by fluorescence microscopy. Bar, 3 µm. (h) Super-resolution microscopy showed that histone H3-yeGFP localized within a hollow circular structure where Npr1 accumulated in Lem2-overexpressing cells. Bar, 1 µm.

